# STRESS: Spatial Transcriptome Resolution Enhancing Method based on the State Space Model

**DOI:** 10.1101/2025.06.25.661624

**Authors:** Xiuyuan Wang, Fei Ye, Xiao Liu

## Abstract

The widespread application of spatial resolved transcriptomics (SRT) has provided a wealth of data for characterizing gene expression patterns within the spatial microenvironments of various tissues. However, the inherent resolution limitations of SRT in most published data restrict deeper biological insights. To overcome this resolution limitation, we propose STRESS, the first deep learning method designed specifically for resolution enhancement tasks using only SRT data. By constructing a 3D structure that integrates spatial location information and gene expression levels, STRESS identifies interaction relationships between different locations and genes, predicts the gene expression profiles in the gaps surrounding each spot, and achieves resolution enhancement based on these predictions. STRESS has been trained and tested on datasets from multiple platforms and tissue types, and its utility in downstream analyses has been validated using independent datasets. We demonstrate that this resolution enhancement facilitates the identification and delineation of spatial domains. STRESS holds significant potential for advancing data mining in spatial transcriptomics.

## Introduction

The development and rapid application of spatially resolved transcriptomics (SRT) technologies provides a powerful tool for researchers to understand gene expression level and tissue heterogeneity in situ^1^. Based on different detection principles, spatially resolved transcriptomic techniques currently have two main branches: technologies based on in situ sequencing or in situ hybridization, and technologies based on NGS and in situ capture^2,3^. The former strategy, including MERFISH^4,5^, osmFISH^6^, CosMx^7^ and seqFISH^8,9^, could achieve single-cell or subcellular resolution by integrating imaging techniques and specially-designed targeted primers to characterize the expression levels of specific genes. However, the number of primers imposes limits on the gene throughput. Conversely, the latter strategy, including Spatial Transcriptomics^10^ (ST), 10X Visium^11^, Stereo-seq^12^ and Slide-seqV2^13^, employs the capture of mRNA in situ at designated spot regions to obtain gene expression levels. The utilization of unique spatial barcodes facilitates the acquisition of spatial localization. This strategy is capable of achieving a higher gene throughput.

Despite the widespread utilization of NGS-based Spatial Transcriptomics techniques, a fundamental issue has emerged: the limited resolution. This challenge can be attributed to two primary reasons. Firstly, the area covered by a single spot is often substantially larger than the size of a single cell. Taking the most popular 10X Visium as an example, the diameter of a single spot is 55 μm, which may cover more than 10 cells, particularly in tissues enriched with immune cells, where it may cover more than 30 cells^14^. The second issue pertains to the low spatial coverage of the spots. In the case of 10X Visium, the spots on the chip are arranged in a hexagonal close-packed pattern, with a distance of 100 μm between each spot. This is substantially larger than the diameter of a single spot, resulting in more than 70% of the chip area undetected. The limited spatial resolution poses challenges for downstream analysis. Due to the discontinuity of the spots and the excessive number of cells covered within a single spot, it will be difficult to characterize the spatial distribution of the gene expression and identify the cell types associated with each spot. Recently, high-resolution spatial transcriptomics technologies, represented by Visium HD^15^, have been developed. This technology reduces the minimum spot size to 2 × 2μm, significantly improving resolution compared to previous generations. However, challenges remain, including low mRNA capture efficiency per spot area and the high costs.

Many methods have been developed to fill the gap of spots and help to assist the downstream missions. BayesSpace^16^ first proposed a spatial resolution enhancing method based on a Bayes model incorporating Markov chain Monte Carlo (MCMC) to estimate model parameters. BayesSpace enhanced spatial resolution by dividing individual spots into 6 and 9 sub-spots on the 10X Visium and ST platforms, respectively, with the limitations hinder comprehensive profiling of genome-wide spatial expression patterns. stLeaen^17^ employs an imputation method that utilizes imaging morphology for the purpose of enhancing the quality of SRT data that is characterized by noise or incompleteness. In alternative methods, such as XFuse^18^ and STIE^19^, deep learning models were employed to enhance the spatial resolution to a sub-spot level or single-cell level. This was achieved by combining spatial transcriptomics with paired scRNA-seq data and high-quality histological image data. Nevertheless, for a considerable number of published studies, such associated data is lacking. To this end, we have developed a model that can achieve resolution enhancement and benefit downstream analysis such as spatial domain identification without reliance on scRNA-seq data or HE images, using only SRT data. In this study, we propose STRESS (Spatial Transcriptomic Resolution Enhancing method based on State Space Model), a structured deep learning-based approach that models gene expression levels in a latent three-dimensional state space integrating spatial coordinates and transcriptional profiles, enabling prediction of spatial gene distributions in a higher scale. Based on this framework, our method achieves a 4-fold enhancement in the resolution for spatial transcriptomic data, thereby increasing the original spatial coverage from 18.3% to 73.2%.

### Overview of STRESS

Recent advances in spatial transcriptomics have enabled genome-wide profiling of gene expression with spatial context. However, the limited spatial resolution remains a major limiting factor, restricting the identification of fine-grained tissue boundaries and spatial domains, as well as the enhancement of gene distribution. Here, we proposed a structured 3D modeling method to enhance the spatial resolution of transcriptomic data, mapping the gene expression matrix to the spatial coordinate matrix, and innovatively constructed a 3D tensor with dimensions *G × H × W,* where *H × W* approximates the spatial layout of spots and *G* the number of genes (Figure 1a). This 3D representation retains both the gene expression landscape and spatial topology, making it amenable to modern deep learning techniques.

**Figure 1:**
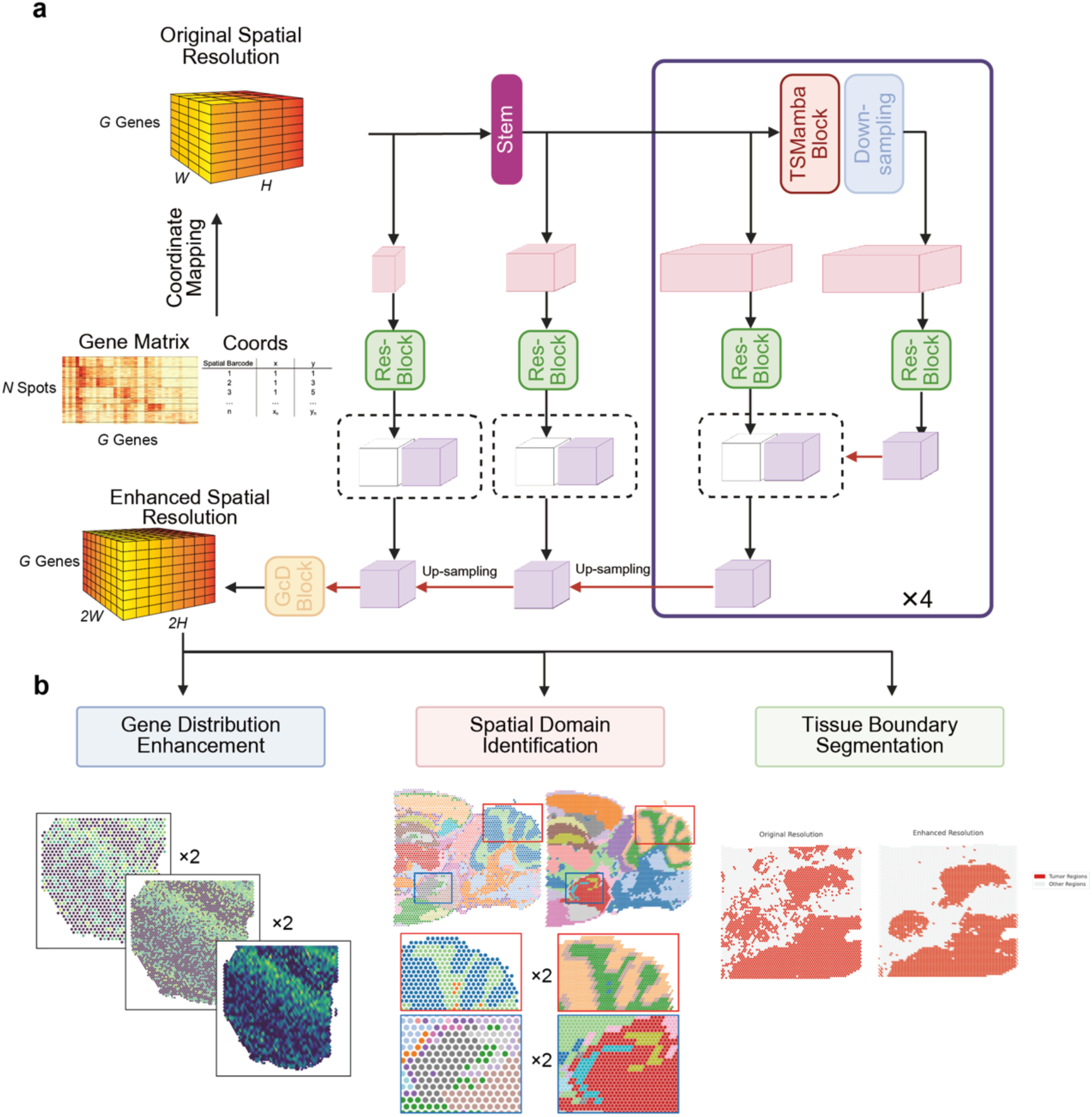
Workflow of STRESS models. **a**,. STRESS takes as input a 3D tensor of dimensions G × H × W, derived from the mapping of gene expression and spatial coordinate information. This input is processed by an encoder composed of a Stem layer followed by four TSMamba blocks with Down-sampling. The resulting features are then passed through a decoder consisting of four Up-sampling modules and a GcD block to enhance spatial resolution. Additionally, Res-Blocks are incorporated to integrate multi-scale encoder features via skip connections. Ultimately, the model outputs a super-resolved 3D tensor with dimensions G × 2H × 2W. **b**, The resolution-enhanced gene expression matrix provided by STRESS is utilized for downstream analyses, including enhanced representation of gene distribution, spatial domain identification in complex tissues, and more precise segmentation of tissue boundaries.

To upscale the spatial resolution, we have, for the first time, developed a dedicated 3D architecture based on the state-space model (STRESS) to accommodate spatial transcriptome properties in spatial transcriptome resolution enhancement task, expanding the boundaries of the state-space model for spatial bioinformatics applications. STRESS follows an encoder-decoder architecture: the encoder consists of a Stem feature projection layer followed by four stacked TSMamba blocks with hierarchical down-sampling, which progressively encode contextual information across spatial and gene-expression dimensions. The decoder adopts a U-net^20^ configuration with symmetric up-sampling, employing Res-Blocks to integrate multi-level encoder features at multiple levels via skip connections. This design preserves fine-grained details while enhancing spatial fidelity. To upscale the 3D tensor, a GcD block is designed as an alternative to the traditional three-directional up-sampling method. This block enhances the resolution of spatial transcriptomic gene expression profiles at the spot level and contributes to smoother tissue region boundaries. STRESS outputs a super-resolved tensor of size *G × 2H × 2W*, effectively doubling the resolution in both spatial dimensions. This architecture provides a scalable and biologically informed solution for reconstructing fine-scale spatial transcriptomic profiles, with strong potential for downstream analytical applications, such as spatial domain identification and tissue boundary segmentation (Figure 1b).

Different from methods such as BayesSpace^16^ that enhance resolution by segmenting capture spots, STRESS directly predicts gene expression in the gaps between original capture spots to achieve resolution enhancement. Consequently, STRESS is more sensitive to the size, distribution density, and arrangement pattern of capture spots. Considering that these characteristics vary across different technological platforms (10X Visium, ST and Stereo-seq), to specifically enhance the resolution of the 10X Visium platform, we employed down-sampling strategy for the training dataset and LOOCV (Leave-One-Out Cross-Validation) training approached. The strategy involves down-sampling the original-resolution dataset as model input while using the original-resolution as Ground Truth, thereby constructing a resolution enhancement mapping function. In order to provide a comprehensive evaluation of STRESS, the performance of the model’s predictions was compared to that of baseline interpolation methods (Nearest Neighbor Interpolation, NNI) as opposed to BayesSpace, with consideration given to the distribution pattern of spots.

Metrics including RMSE, PGCC, PSNR, and SSIM were used to assess the similarity between the predicted high-resolution spatial transcriptional profiles and the true high-resolution spatial transcription pips (see Methods).

### STRESS enhanced spatial resolution of human DLPFC dataset

Firstly, we collected LIBD human dorsolateral prefrontal cortex (DLPFC)^11^ dataset generated from 10X Visium platform as the training set. For each sample, we selected the top 2048 highly variable genes (HVGs) and constructed a three-dimension matrix incorporating spatial coordinates and gene expression. The results in DLPFC dataset show that STRESS’s prediction is significantly higher than NNI in RMSE, PGCC, PSNR and SSIM metrics (Figure 2a). In particular, the average value of PGCC, PSNR, and SSIM improved by 25.6%, 225.71%, and 7.95% compared to NNI. The average value of RMSE decreased by 8.66% compared to NNI. The predictions indicate that the STRESS method is more accurate, and the resolution-enhanced prediction results are very similar to the actual situation.

**Figure 2:**
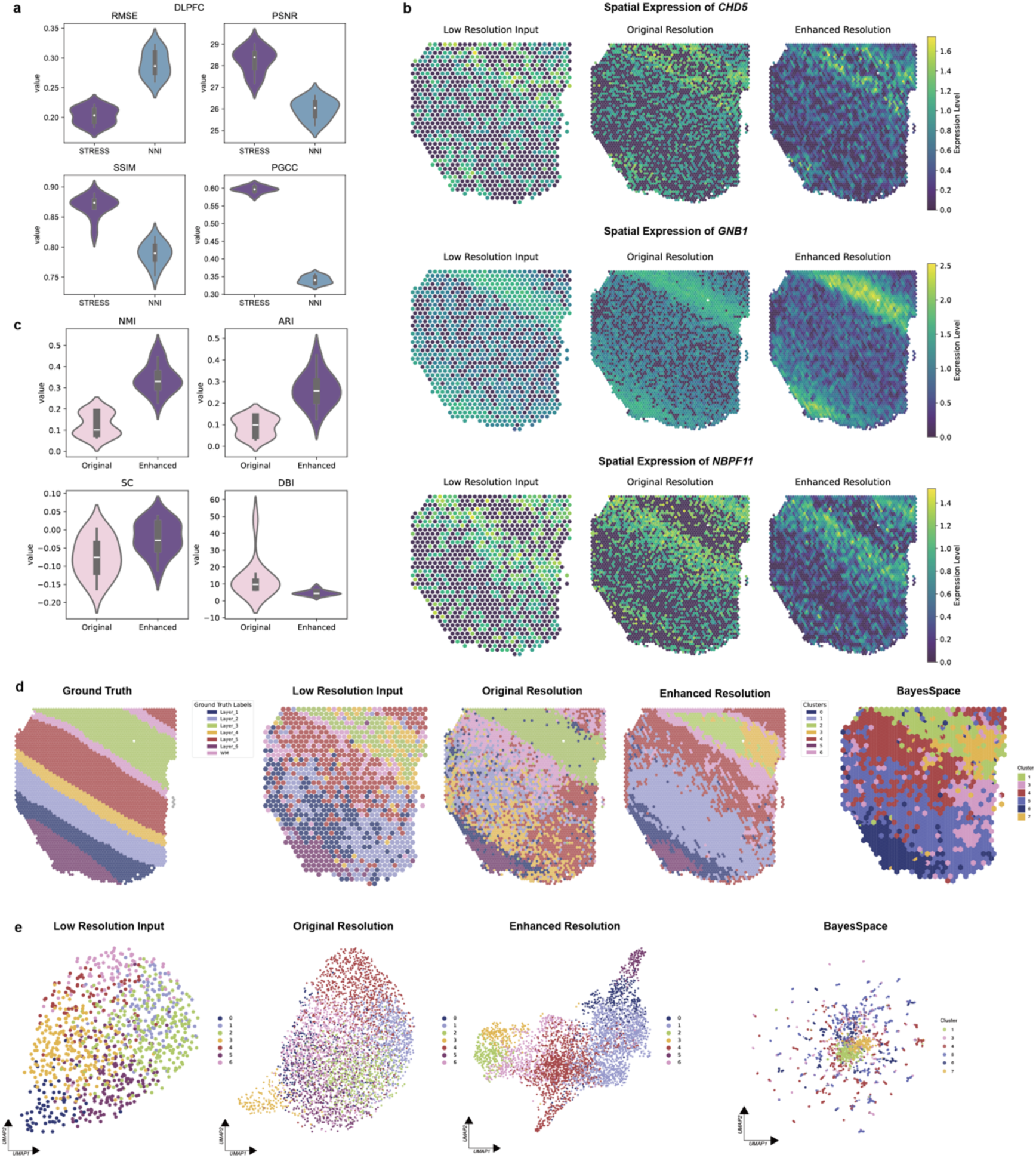
STRESS provides accurate resolution-enhanced gene distribution predictions that facilitate clustering implementation. **a**, Violin plots of evaluation metrics between STRESS and baseline (NNI) results in DLPFC dataset. RMSE, Root Mean Square Error; PSNR, Peak Signal-to-Noise Ratio; SSIM, Structural Similarity Index; PGCC, per-gene expression correlation coefficient (Details see the Methods). **b**, Spatial expression patterns for different genes (*CHD5*, *GNB1*, and *NBPF11*) across low resolution input, original resolution (Ground Truth), and STRESS-enhanced resolution prediction. **c**, Violin plots of clustering metrics between low resolution input and enhanced resolution prediction. NMI, Normalized Mutual Information; ARI, Adjusted Rand index; SC, Silhouette coefficient; DBI, Davies-Bouldin Index. **d**, Manual annotation of slice 151507 in DLPFC dataset, spatial domain identification results based on low resolution input, original resolution Ground Truth and STRESS-enhanced resolution prediction using same clustering method and same parameters, and spatial domain identification results of low-resolution input based on BayesSpace. **e**, Latent representations of slice 151507 in DLPFC dataset obtained by performing UMAP dimensionality reduction on gene expression matrices of low-resolution input, original resolution Ground Truth, enhanced resolution prediction, and BayesSpace result, with different colors representing previously identified domain labels.

Gene-specific validation using spatially variable genes from the DLPFC dataset (Figure 2b) revealed that enhanced predictions maintained regional enrichment patterns observed in ground truth annotations. For example, *GNB1* showed layer-specific enrichment in cluster 2 (corresponding to annotated Layer 1), while *CHD5* and *NBPF11* exhibited precise sequential localization in clusters 6 and 4 (matching Layers 2 and 3, respectively), with expression gradients mirroring known cortical laminar organization^21^. Another resolution enhancement algorithm, BayesSpace, was also applied to the low-resolution data. In comparison with the gene expression distribution that has been enhanced by BayesSpace (Supplementary Figure 1), the results obtained from STRESS are more closely aligned with those that have been obtained at original-resolution. Furthermore, it exhibits a notable boost compared to other regions, indicating that the model more accurately identifies the gene’s expression pattern.

The aforementioned performance evaluations are all based on the accuracy of the spatial gene expression matrix. In fact, the performance of the model’s output results in downstream analyses is also crucial for assessing the usability of the model. Downstream analytical utility was assessed through unsupervised clustering of the input and output of STRESS. Evaluation metrics included Normalized Mutual Information (NMI) and Adjusted Rand Index (ARI) for clustering accuracy, supplemented by Silhouette Coefficient (SC) and Davies-Bouldin Index (DBI) for spatial consistency (see Methods). Based on the same clustering method with the same parameters, the resolution enhanced ST expression shows better clustering effect and regional consistency than the low-resolution input (Figure 2c). The clustering visualization results for one of the samples demonstrate the accuracy of the enhanced-resolution predictions (Figure 2d). The spatial domain segmentation results have clearer boundaries compared to low resolution input, while the prediction results can better discriminate between similar regions in close proximity compared to original-resolution. Comparison with BayesSpace method demonstrated that STRESS-enhanced resolution predictions generated outputs more closely resembling the actual spot distribution patterns (Figure 2d). Unlike BayesSpace that assigns each spot to neighboring similar spots, our predictions successfully identified and reconstructed the distinct spatial distribution of all domains while better preserving the true spatial organization. Furthermore, UMAP of gene expression (Figure 2e) revealed that the enhanced resolution predictions better capture transcriptional heterogeneity, leading to superior clustering performance. In contrast, the BayesSpace tended to produce similar gene predictions. For the majority of samples (Supplementary Figure 2, 3), the predicted clustering performance is on par with or surpasses that of the Ground Truth before down-sampling. This outcome suggests that the STRESS model can achieve superior clustering performance using reduced input (only 1/4 of the original spots), by learning the relationships both between genes and between genes and their spatial positions.

### STRESS performance validation on independent dataset with Ground Truth

The performance of the STRESS model was then tested on independent datasets with Ground Truth using the STRESS-Visium parameters obtained from previous training methods. It has been demonstrated that STRESS performs well in the “down-sampled resolution to original resolution” mapping scenario (64×64 to 128×128). However, in practical applications, the “original resolution to enhanced resolution” mapping relationship (128×128 to 256×256) is frequently encountered. Consequently, independent validation is required under this mapping paradigm. Emerging methods such as Visium HD and Slide-seq, with their higher resolutions, can be utilized to create the required mapping relationships (128×128 as input and 256×256 as Ground Truth). Due to the limited sample sizes in these datasets being insufficient for training, a selection of six Visium HD samples (see Methods) and one Slide-seq V2^13^ sample was made for independent validation dataset, with the objective of demonstrating STRESS’s stability and robustness. The Visium HD dataset integrates data from 8-micron bin mode and 16-micron bin mode, using 128×128 data as input and 256×256 data as Ground Truth (see Methods). The results demonstrate that STRESS’s predictions on the Visium HD dataset manifest a notable congruence with the Ground Truth, exhibiting a mean PSNR of 41.01, a mean SSIM of 0.9653, a mean RMSE of 0.1694, and a mean PCC reaching 0.6710 (Figure 3a). This performance exceeds that of the training set (mean PSNR of 28.24, mean SSIM of 0.8682, mean RMSE of 0.2028 and mean PCC of 0.5966 in the training set, Figure 2a). The findings indicate that STRESS is capable of maintaining consistent performance across a range of resolution mapping relationships and can effectively facilitate resolution enhancement for particular applications.

**Figure 3:**
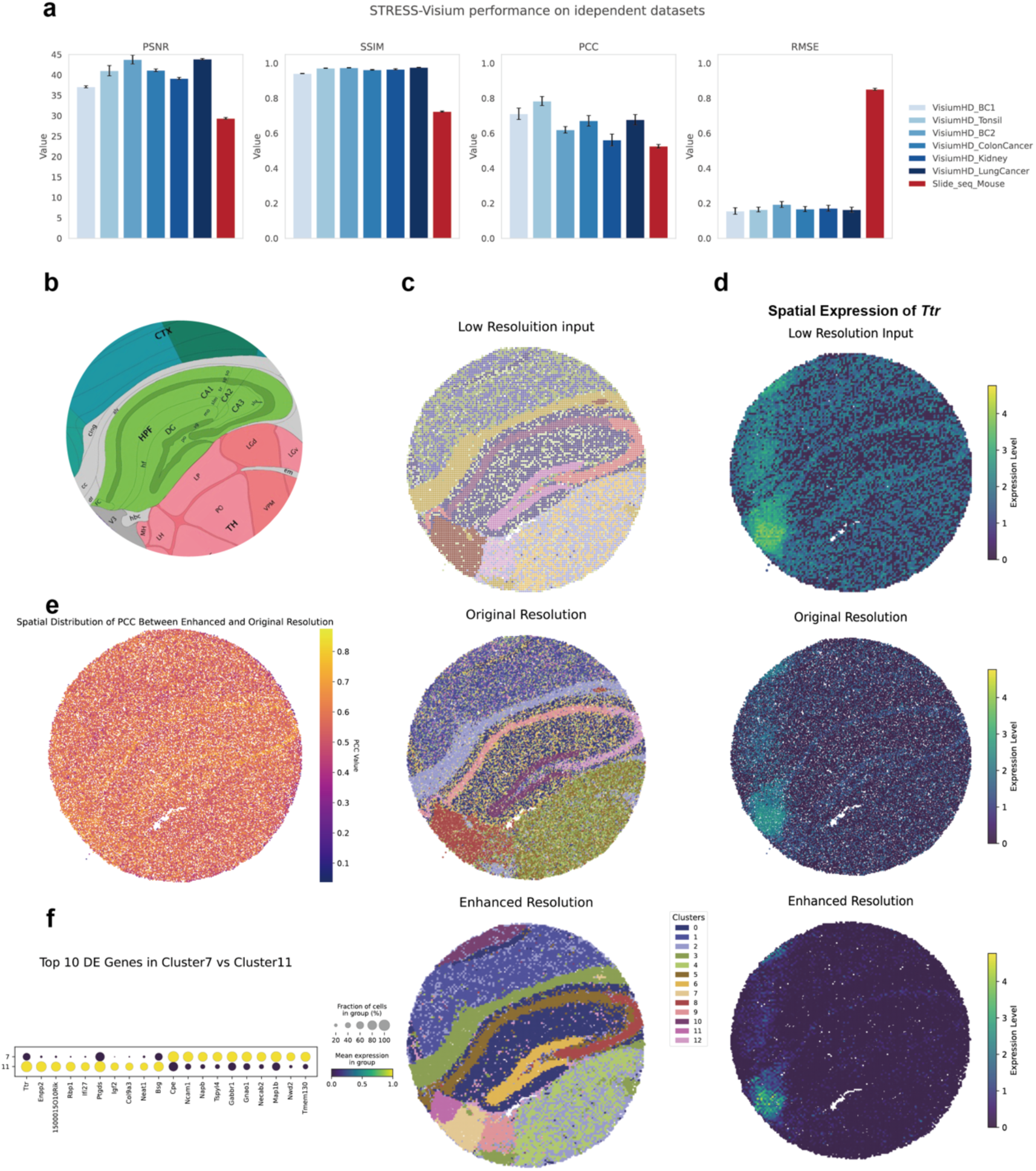
Independent validation of STRESS on datasets with ground truth. **a**, Histogram of STRESS-Visium model performance on different samples of independent datasets (Visium HD and Slide-seq V2). RMSE, Root Mean Square Error; PSNR, Peak Signal-to-Noise Ratio; SSIM, Structural Similarity Index; PCC, per-gene expression correlation coefficient. **b**, The Allen Brain Atlas^23,38^ annotated layer structure of mouse hippocampus. CTX, Cerebral cortex; HPF, Hippocampal formation; cing, cingulum bundle; alv, alveus; CA1slm, Field CA1, stratum lacunosum-moleculare; DG-mo, Dentate gyrus, molecular layer; sg, granule cell layer; po, polumorph layer; FC, Fasciola cinerea; V3, third ventricle; hbc, habenular commissure; MH, Medial habenula; LH, Lateral habenula; LP, Lateral posterior nucleus of the thalamus; LGd, Dorsal part of the lateral geniculate complex; LP, Lateral posterior nucleus of the thalamus; PO, Posterior complex of the thalamus. **c**, Spatial domain identification results in mouse hippocampus dataset based on low resolution input, original resolution Ground Truth and STRESS-enhanced resolution prediction using same clustering method and same parameters. **d**, Spatial expression patterns for *Ttr* gene across low resoluition input, original resolution (Ground Truth), and STRESS-enhanced resolution prediction. **e**, Spatial distribution of Pearson Correlation Coefficient (PCC) of gene expression vector in each spot between original resolution and enhanced resolution prediction. **f**, Top 10 differentially expressed genes between cluster 7 and cluster 11.

The mouse hippocampus Slide-seq V2^13^ dataset contains 51,367 spots with the size of 10 micrometers, approaching single-cell resolution, and more than 20,000 genes. In order to provide further validation of the model’s generalizability and robustness, we take the Slide-seq dataset to simulate 10X Visium distribution pattern and used it as an independent testing dataset to test STRESS-Visium model. We filtered the genes and selected top 2048 HVGs. The originally discrete spots of Slide-seq are covered by 128×128/256×256 regular grids, thereby generating simulated 3D spatial gene expression matrix with different resolutions. STRESS utilized 128×128-resolution data as the input layer and provided a 256×256-resolution prediction. The STRESS-Visium prediction results based on independent Slide-seq V2 dataset still show a high level of similarity with the Ground Truth, achieving a mean PCC of 0.5262. Additionally, the mean PSNR, mean SSIM and mean RMSE are 29.36, 0.7245 and 0.8513, respectively (Figure 3a). To further validate the effectiveness of STRESS’s resolution enhancement in downstream analyses, we selected the prediction results from the DLPFC-trained STRESS model for downstream analysis. We observed strong spatial distribution similarity between the predicted gene matrix (at enhanced resolution) and ground truth (at original resolution) (Figure 3e, median PCC value = 0.56). In Figure 3c, we compared the clustering results generated from low-resolution input, original-resolution Ground Truth and enhanced-resolution prediction using the same clustering method and same parameters. It was evident that a clear layered structure of the mouse hippocampus was observed in the enhanced-resolution prediction, which exhibited a high degree of consistency with the original-resolution Ground Truth. In comparison with the low-resolution input, the clustering result generated from STRESS was able to identify more specific regions, such as V3 (third ventricle) and hbc (habenular commissure), which were consistent with the Allen Brain reference atlas^22^ annotations. Meanwhile, the enhanced-resolution prediction exhibited higher spatial heterogeneity across different layers in comparison with the original-resolution Ground Truth (Figure 3c, Supplementary Figure 4). For instance, the predictions distinguished the difference between the sg (dentate gyrus, granule cell layer) and LH (Lateral Habenula) region (Cluster 7 and Cluster 11 in Figure 3c), which was identified as a single region in the original-resolution Ground Truth. Based on predicted gene matrix and clustering results, we conducted the differential expression analysis and identified the top enriched genes in different regions (Figure 3f). Consistent with previous conclusions, we observed a high expression level of *Ttr* gene (Figure 3d), which is a V3 choroid plexus marker gene involved in thyroxine transport and cerebrospinal fluid secretion^23,24^. The above conclusions demonstrate that STRESS can effectively enhance spatial resolution and better identify heterogeneity among spatial domains.

In order to provide further evidence that the STRESS architecture’s resolution enhancement effect is applicable to different platforms and tissues, the same strategy was adopted as previously, with models being trained using the HER2ST^25^ dataset from the ST platform and the MOSTA^12^ dataset from the Stereo-seq platform. The resulting STRESS-ST and STRESS-Stereo models were then validated on independent VisiumHD and Slide-seq datasets. The findings of the training process demonstrate that the predictions derived from STRESS-ST and STRESS-Stereo models demonstrate a high degree of similarity with Ground Truth (Supplementary Figure 5, 6), whilst concurrently exhibiting optimal performance metrics in independent dataset testing (Supplementary Figure 7). The results obtained from this study suggest that STRESS is an effective method for learning gene interaction relationships between spots and provides accurate, resolution-enhanced predictions based on a three-dimensional state-space model. In order to maintain platform consistency, it is imperative that all subsequent validations are based on the STRESS-Visium model parameters.

### STRESS model helps identify clusters with subtle heterogeneity

Next, to further validate the benefit of STRESS-predicted enhanced-resolution gene expression in downstream analysis, we selected an additional independent mouse brain Visium dataset^26–29,30^, encompassing four samples that represented 2 anterior and 2 posterior sections adjacent in mouse brain, respectively. Here, a posterior section was selected for further analysis (Section 1 of the Posterior, HE image in Figure 4a). A comparison was made of the clustering results obtained using Leiden clustering method^31^ on both the original-resolution and STRESS enhanced-resolution data, as well as the SpaGCN result on the original-resolution data (Figure 4b). Initially, the number of clusters was set at 8, as determined by the elbow method in K-Means clustering (Supplementary Figure 8). The results demonstrate the efficacy of all clustering methods in identifying the spatial structures of the mouse brain. However, when compared to original-resolution + Leiden, both STRESS enhanced-resolution + Leiden and SpaGCN + original-resolution yield clustering results with stronger spatial continuity and integrity. Furthermore, the STRESS enhanced-resolution + Leiden approach successfully identified the HPF region, which is in accordance with the Allen Brain Institute reference atlas (see red box in Figure 4b and Supplementary Figure 9). In order to conduct a more in-depth investigation into STRESS’s performance in identifying finer tissue structures, the cluster number was increased to 20. In accordance with the findings of previous results, the clustering results at both the original and enhanced resolutions were able to identify more detailed brain structures (Supplementary Figure 10). However, the Louvain clustering result of STRESS enhanced-resolution data revealed a complete granule cell layer and layered organization in the Cerebellar Cortex region, which aligns with HE images and annotations from the Allen Brain Institute reference atlas. The results presented herein demonstrate that STRESS enhanced-resolution data facilitates the discovery of clusters within complex structures.

**Figure 4:**
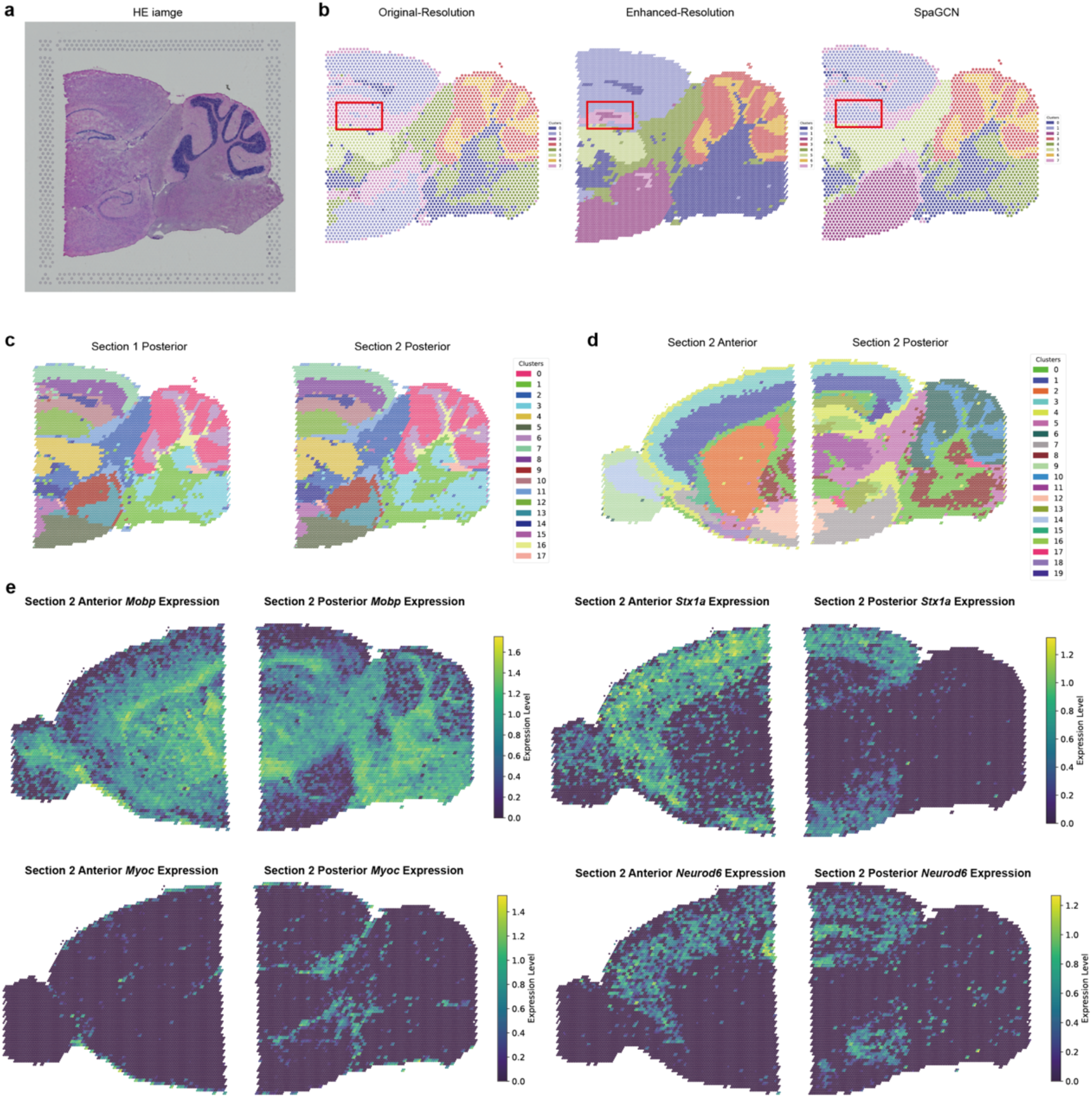
STRESS helps with spatial domains identification in complex mouse brain tissues **a**, Histology image of mouse brain posterior tissue section 1^26^. **b**, Spatial domain identification results in mouse brain posterior tissue section 1, generated using Leiden clustering method at original resolution, Leiden clustering method at STRESS-enhanced resolution and SpaGCN clustering method at original resolution. Red box, HPF region corresponding to the Allen Brain Institute reference atlas. **c**, Integration of spatial domain identification results from two consecutive mouse brain posterior tissue, generated at STRESS-enhanced resolution. **d**, Integration of spatial domain identification results from two adjacent mouse brain tissue, generated at STRESS-enhanced resolution. **e**, Distribution of cluster-specific genes expression (*Mobp*, *Stx1a*, *Myoc*, and *Neurod6*) across two adjacent samples, anterior section 2 and posterior section 2.

The results mentioned above were derived from individual sample analysis. However, our focus is equally weighted towards the consideration of spatial regional heterogeneity across consecutive or adjacent sections. Consequently, we subsequently performed resolution enhancement on consecutive sections from the same region and adjacent sections at the same depth within the dataset. STRESS was applied to each sample individually, followed by integrated clustering analysis of the enhanced-resolution data. As illustrated in Figure 4c, the clustering results of two continuous sections from the same region of the mouse posterior demonstrate a high degree of similarity, which is consistent with HE images. Figure 4d presents the clustering results of two adjacent sections from the anterior and posterior mouse brain in Section 2. Given the anatomical proximity of these two sections, it was observed that there were numerous clusters spanning both sections, which could be seamlessly stitched together at the junction. Based on the clustering results, we identified differentially expressed genes in these cross-section clusters (Supplementary Figure 11) and visualized their spatial distribution patterns (Figure 4e and Supplementary Figure 12). According to the analyses of differentially expressed genes, cluster 0 highly expressed genes that were previously found to be up-regulated in Fiber Tracts. The distribution of cluster 0 was found to be consistent with Fiber Tracts according to manual labels, usually at the junctions among different regions. More clusters were also found to distribute consecutively between the two samples, such as cluster 3 (highly expressing Cerebral cortex related genes), cluster 4 and cluster 13. The results of the clustering process and gene expression distributions demonstrate that STRESS exhibits strong stability and robustness in performance across different tissue slices and emphasize the potential of STRESS in the mission of enhanced-resolution structural construction of tissue samples.

### STRESS model helps with tissue boundary segmentation in tumor microenvironment

In the tumor microenvironment, tissue boundaries segmentation is crucial for studying interactions between tumor and normal tissue. To validate STRESS’s performance in boundary identification, we selected a human breast cancer sample^32^ on 10X Visium platform as independent dataset for testing. The sample was collected from an invasive ductal carcinoma (IDC) patient who has been detected as estrogen receptor positive (ER+), progesterone receptor negative (PR-) and human epidermal growth factor receptor 2 positive (HER2+). Based on previous processing pipelines, the clustering results at original resolution and STRESS-Visium enhanced-resolution are shown in Figure 5a. We observe that the latent representation of clustering result based on STRESS enhanced-resolution has the capacity to more effectively identify gene expression patterns across different regions compared to original-resolution (Figure 5b), particularly in areas exhibiting tumor heterogeneity.

**Figure 5:**
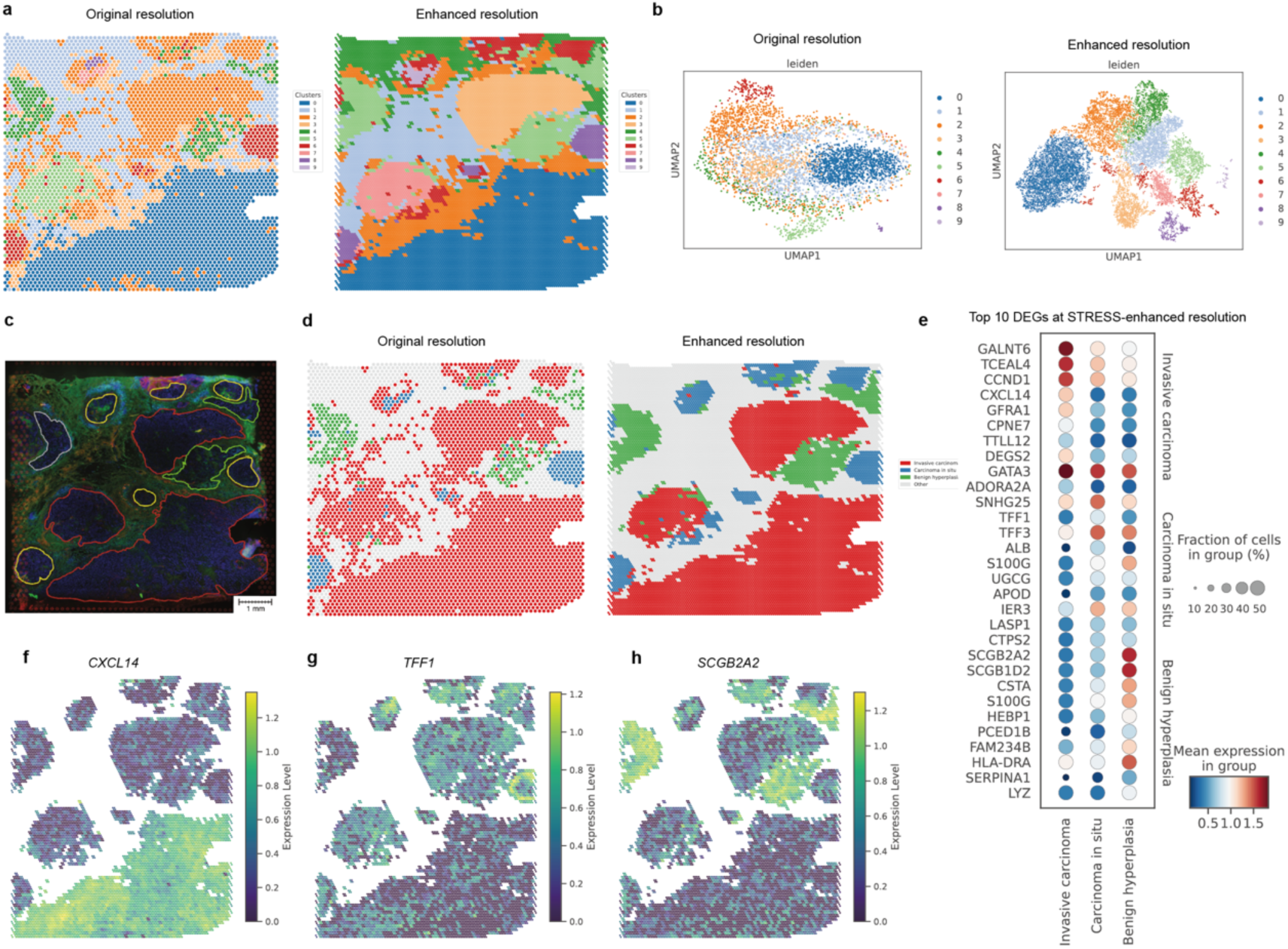
STRESS prediction helps with domain boundary segmentation in Tumor Microenvironment. **a**, Spatial domain identification results in IDC sample, generated using Leiden clustering method at original resolution and STRESS-enhanced resolution. **b**, Latent representations of IDC sample obtained by performing UMAP dimensionality reduction on gene expression matrices of original resolution input and STRESS-enhanced resolution prediction, with different colors representing spatial domain labels. c, Immunofluorescent imaging of the IDC sample with manual annotations, cite from *Zhao E et al.*^16^ Invasive carcinoma regions are outlined in red, carcinoma in situ regions are outlined in yellow, benign hyperplasia region is outlined in green and the unclassified tumor region is outlined in gray. **d**, Distribution of Invasive carcinoma region, Carcinoma in situ region and Benign hyperplasia region at original and STRESS-enhanced resolutions. **e**, Top 10 differential expression genes among Invasive carcinoma region, Carcinoma in situ region and Benign hyperplasia region at STRESS-enhanced resolution. **f-h**, Spatial distribution of *CXCL14*, *TFF1* and *SCGB2A2* genes expression.

In previous studies by *Zhao E et al.*^16^, the identification of distinct regions of invasive carcinoma, carcinoma in situ, and benign hyperplasia within the tissue was achieved through the use of pathological images by pathologists (Figure 5c). According to the results of the gene expression clustering, these regions corresponded to different clusters at both original and enhanced resolutions: at original resolution, invasive carcinoma corresponded to cluster 0, 3, and 7; carcinoma in situ to cluster 6, 8, and 9; and benign hyperplasia to cluster 5. At STRESS enhanced-resolution, invasive carcinoma corresponded to cluster 0, 5, and 2; carcinoma in situ to cluster 6 and 7; and benign hyperplasia to cluster 4. As demonstrated in Figure 5d, in comparison with the original resolution, the invasive carcinoma region identified at STRESS-enhanced resolution was distinctly more clearly delineated, with the average Silhouette score rising from 0.0613 to 0.5119 (Supplementary Figure 13). Furthermore, the enhanced resolution enabled accurate delineation of the boundaries of carcinoma in situ and benign hyperplasia, consistent with expert annotations (Figure 5c). In order to ascertain the cellular phenotype and function of these three regions, differential gene expression analysis was performed on the STRESS-enhanced resolution data (Figure 5e). In the invasive carcinoma region, the elevated expression of the chemokine, *CXCL14* (Figure 5f), is a common hallmark of cancer associated fibroblast (CAF), while the upregulation of genes such as cyclin D1(*CCND1*) is frequently observed in ER+ invasive breast cancer (Supplementary Figure 14). Genes such as *COL1A1* and *FN1* appear to be indicative of the invasive and motile characteristics of this region (Supplementary Figure 14), which is consistent with the features observed in invasive carcinoma. Conversely, in the carcinoma in situ region, the upregulation of *TFF1/TFF3* genes is frequently observed (Figure 5g and Supplementary Figure 14), reflecting the early stages of carcinogenesis. The benign hyperplasia region exhibited differential expression of the *SCGB2A2* gene (Figure 5h), which is associated with benign breast lesions, along with increased expression of immune-related genes, including *HLA-DRA* and *LYZ* (Supplementary Figure 14). The findings outlined above serve to underscore the potential of STRESS-enhanced resolution results in the analysis of the tumor microenvironment, helping to more precisely define tumor regions and boundaries.

## Discussion

In this study, we presented STRESS, the first deep learning framework that leverages spatial transcriptomics data to construct a 3D latent architecture for resolution enhancement without the reliance of paired scRNA-seq data and HE image. STRESS is specially designed for Visium datasets; however, evidence has been provided demonstrating its capacity to be trained on multi-platform (including ST and Stereo-seq), multi-tissue-type datasets to achieve comparable performance. Our results demonstrate that STRESS maintains stable performance when applied to Visium data simulated from high-resolution datasets such as VisiumHD and Slide-seqV2, successfully enhancing resolution by a factor of 4 compared to the input. Furthermore, we validated STRESS’s performance on multiple independent datasets and demonstrated that this resolution improvement benefits downstream missions, including spatial domain identification and tumor boundary delineation. Application to a mouse brain dataset reveals STRESS’s ability to preserve biological continuity across adjacent and continuous slices, which shows the potential of STRESS on tissue panoramic reconstruction. In another human breast cancer dataset, as a result of the resolution-enhanced results from STRESS, clearer tumor boundaries were identified, and a comparison was made of the gene expression heterogeneity of tumors at different locations—which could be challenging to detect at the original resolution. Additionally, STRESS’s high-resolution spatial domain identification achieved performance comparable to the resent spatial domain identification algorithms, such as SpaGCN. Notably, STRESS accomplished this using only gene expression and spatial location information, without relying on additional data such as H&E staining images or paired single-cell transcriptomics, which are commonly utilized by other algorithms. In summary, a wholly novel, auxiliary information-free model framework STRESS is proposed for achieving resolution enhancement of Visium spatial transcriptomic data, thus providing a powerful tool for spatial transcriptomics data mining.

STRESS is capable of high-throughput processing of the full gene expression profiles in spatial transcriptomics data. We ensured that STRESS can be easily deployed and used on local servers. STRESS requires approximately 4 seconds to process a single sample with 2048 genes during inference, evaluated on a server equipped with an AMD EPYC 9754 CPU and a single NVIDIA H800 GPU. Additionally, to improve the model’s performance on new datasets, we also provide a method for fine-tuning parameters through re-training on new datasets.

While STRESS shows robust performance for resolution enhancement, there exists challenges and limitations. One limitation of STRESS is that its predictions are entirely based on the spatial relationships and interactions among the input gene sets. Technical artifacts such as mRNA drifts and the sparse expression of some specific genes may lead to false-positive imputation. Constrained by computational resources, the top 1024 highly variable genes were selected as input during training. The incorporation of additional genes may result in enhanced model performance. Although STRESS performs well in tasks such as spatial domain identification based on gene expression, predictions for individual genes would require the integration of additional modalities, such as H&E staining images or paired scRNA-seq datasets, into the model. We will address above limitations into our further work by adding gene label as model input and integrating imaging data as an additional module into our model.

## Methods

### Dataset collection and preprocessing

STRESS was trained and tested on three datasets: **Human dorsolateral prefrontal cortex dataset** (DLPFC) generated from 10X Visium platform. **Human HER2-positive breast cancer dataset** (HER2ST) generated from Spatial Transcriptomics (ST) platform. **Mouse Organogenesis Spatiotemporal Transcriptomic Atlas** (MOSTA) generated from Stereo-seq platform.

More downstream analysis was validated on independent datasets: **Mouse hippocampus dataset** generated from Slide-seqV2 platform was used to simulate 10X Visium data. **Mouse brain serial section dataset** generated from 10X Visium platform. **Human invasive ductal carcinoma dataset** generated from 10X Visium platform. **Visium HD dataset** generated from 10X Visium HD platform.

**Preprocessing method**: Normalization and principal components analysis (PCA) dimensionality reduction step was applied on each sample using *scanpy* Python package (version 1.10.2). Highly variable genes (HVGs) were identified using ‘highly_variable_genes’ function in *scanpy* Python package (version 1.10.2) and top 2048 HVGs were selected as input of STRESS.

### Design of STRESS Problem definition

For a spatial transcriptome sample, we first convert it into a 3D matrix *X* ∈ *R*^1xGxHxW^ based on its coordinates and corresponding gene expression profile, which represent that the number of spots is *H* × *W* and there are G genes for each spot. Our goal is to up-scale each spot *N* folds with keeping the number of genes constant by using deep learning and final get a super-resolution matrix *Y* ∈ *R*^1xGxNWxNH^.

**STRESS** is a encoder-decoder structure. For the encoder, there are five stage to extract the shallow and deep information. The shallow information prefers to learning the contouring information and the deep stage prefers to learning the detail information.

### Tri-orientated Spatial Mamba Block (TSMamba)

Modeling local and global feature and exploring the interaction among spots and genes are critically important for enhancing spatial transcriptomic gene expression profile. Conventional spatial transcriptome super-resolution methods often use 2-dimensional deep learning architectures, and they usually only take into account the interrelationships between spots, and are lacking in the extraction of gene-to-gene feature information. We thus consider for the first time a 3D deep learning architecture to explore the spatial transcriptomic data enhancement. In general, CNN-based and Transformer-based method are widely used as a feature extractor. However, CNN-based architecture can only extract local feature information within a specific receptive field limited by the fix size of convolutional kernel and Transformer-based architecture can learn feature information for all spots and genes but it incurs a significant computational burden especially for the big input matrix. Based on this, we design 3D mamba-based encoder^33^, originates from state space models^34^, to enable multi-scale feature modeling while maintains a high efficiency during training and inference.

The encoder contains of a Stem feature projection layer, four stacked TSMamba blocks with down-sampling. For the Stem layer, we use a 3D depth-wise convolution layer with a kernel size of 7 × 7 × 7, a padding of 3 × 3 × 3, a stride of 2 × 2 × 2, and channel number of 48. Given a spatial transcriptome matrix *I* ∈ *R*^CxGxWxH^, where *C* represent the number of input channels, the initial feature 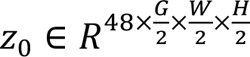 is extracted by the Stem layer. Then, the *z*_0_ is fed through four TSMamba and corresponding down-sampling. For the *t*^th^

TSMamba block, it can be definite as

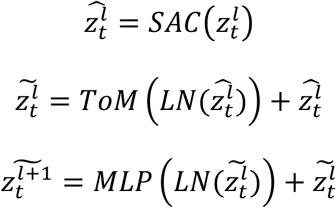

Where the SAC and ToM denote the proposed gated spatial convolution module and tri-orientated Mamba module, respectively, which will be discussed next. *l* ∈ {0, 1, …, *N*_m-1_}, LN denotes the layer normalization, and MLP denote multiple layers perceptions.

### Spatial auxiliary convolution (SAC)

The mamba layers learning feature representation by flattening the 3D feature into 1D sequence, which loses the contextual information from spot to spot and fails to learn the differences between the whole different organization regions of the input features. Hence, we design a Spatial auxiliary convolution (SAC) module. The input 3D features *z* are respectively fed into a *CB*_1_, *CB*_2_, and *CB*_3_, with the convolution kernel sizes of *CB*_1_ and *CB*_2_ being 3 × 3 × 3 and the *CB*_3_ being 1 × 1 × 1, then the two features are added pixel-by-pixel to complement the various scales of information extracted. Finally, *CB*_)_ is used to further fuse the feature, while a residual connection is utilized to reuse the input feature.

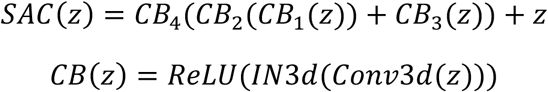

Where *CB*_i_ denote spatial convolution block, *ReLU* is an activation function, and *IN*3*d* is 3D instance normalization, and *Conv3d* denote 3D convolution operation.

### Tri-orientated mamba (ToM)

Traditional mamba layers are all extract feature information from two directions, that is the width and height for a fix pixel image or fix spot spatial transcriptome gene expression profile, which lacks the ability to learn information about the differences of different genes in a multi-spot neighborhood. Hence, we thus design a tri-orientated mamba module that compute the feature representations from three directions, that is gene, width, and height for a fix spot spatial transcriptome gene expression profile, to further extract spatial high dimensional information. We flatten the 3D feature into three sequences and input mamba layer to extract global feature information and obtain fused 3D feature maps.

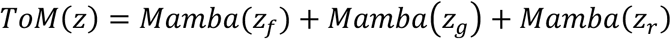

Where *Mamba* is the mamba layer to extract the global information of each sequence, *z*_f_ denotes the forward directions, *z*_r_ denotes reverse direction, and *z*_s_ denotes inter-gene direction.

### Up-sampling

Due to the large amount of spatial transcriptome gene expression profiling data, we designed a multi-stage encoder to downscale the features in order to ensure that the computational burden is reduced while exploring gene-to-gene and point-to-point feature characterization more comprehensively. Therefore, in this section, we design a decoder, which is a similar procedure as the encoder, for feature recovery. For the decoder, there are also Up-Block to reconstruct feature maps coming from the encoder step by step. Up-Block follows many precious studies ^35–37^, which a CNN-based blocks with a residual connection for reconstructing the feature maps coming from the encoder.

### Gene-constrained dilatation (GcD)

Finally, we propose for the first time a novel up-sample method, that is Gene-constrained dilatation, to enhance spatial resolution for four times in the spot level with the genetic dimension constrained. GcD employs a new paradigm of gene constraints to replace the traditional 3D three-directional up-sampling method to achieve improved resolution of spatial transcriptome gene expression profiles at the spot level, which smoothes tissue region boundaries, discovers unknown new cell types, and depicts tissue microenvironments. The 3D reconstructed feature z is fed into three convolution blocks (a 3D convolution layer with the convolution kernel sizes being 3×3×3, a LeakyReLU), then the features are up-sampled by using the proposed GcD. Given a reconstructed low-resolution data, the sampling factor (*f*_x_, *f*_y_, *f*_g_) of high-resolution in three direction is 1, 2, 2, respectively. The gene expression values *f*(*x*, *y*, *g*) with coordinates (*x*, *y*, *g*) in the predicted high-resolution matrix can be definite as

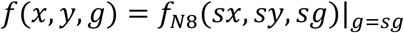

Where *f*_N8_(*sx*, *sy*, *sg*) denote a trilinear interpolation function on the coordinate (*sx*, *sy*, *sg*) of the original image corresponding to the target image at that point (*x*, *y*, *g*). Given the coordination of low-resolution data is (*sx*, *sy*), its maximum coordinate (*sx*_1_, *sy*_1_) and minimum coordinate (*sx*_0_, *sy*_0_) are 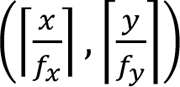 and 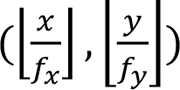 respectively, and their corresponding deconvolutional gene expression are 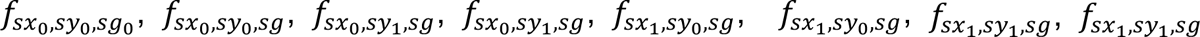. The sampling procedures are followed by three steps. We first interpolate in the *g*-direction, which are

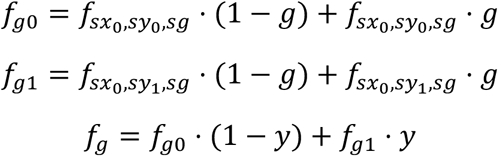

Next, we interpolate in the *y*-direction, it is calculated by

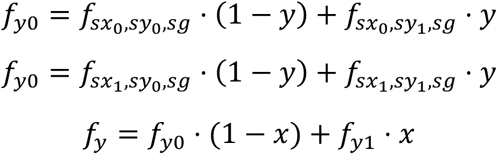

Next, we interpolate in the *x*-direction and obtain final the high-resolution spatial transcriptomic gene expression profile.

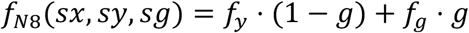

### Training dataset construction

STRESS adopts a method that involves down-sampling the original resolution dataset as model input while using the original resolution as Ground Truth. Considering the size, distribution density, and arrangement pattern of capture spots differ in three different training datasets. We employed a unified down-sampling strategy: based on the parity of spot coordinates, we partitioned every four adjacent spots into four distinct down-sampled subgraphs, thereby generating low-resolution inputs at 1/4 of the original size. Due to variations in dataset dimensions, the high-low resolution correspondence in the three training sets were: HER2ST, 32×32-16×16; DLPFC, 128×128-64×64; Stereo-seq, 128×128-64×64.

### Evaluation metrics RMSE

Root Mean Square Error (RMSE) measures the degree of deviation between the predicted value and the true value, and is calculated as the square root of the sum of the squares of the differences between the predicted value and the true value. The value of RMSE ranges from 0 to positive infinity, and the smaller the value is, the smaller is the prediction error and the better is the prediction ability of the model.

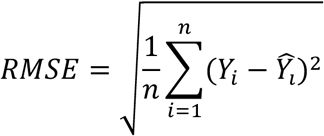

Where *Y*_i_ is the value of the ground truth *i,ŷ_i_* is the value of the predicted *i*, *n* is the total number of predicted values.

### per-gene expression correlation coefficient (PGCC)

To measure the correlation between predicted and true values, we propose a new correlation evaluation metric, the per-gene expression correlation coefficient (PGCC), to measure the correlation between the predicted value and the true value for each gene in each SPOT, which reflects the overall degree of correlation between the observed variable and the true variable, and the closer the value is to 1, the better.

For a given predicted gene expression matrix *X* ∈ *R*^1×G×W×H^ and its ground truth *Y* ∈ *R*^1×G×W×H^, we first flatten them into *x* ∈ *R*^[G×W×H]×1^ and *y* ∈ *R*^[G×W×H]×1^, G, W, and H are number of gene, width, and height, respectively. The correlation between *x* and *y* is then calculated by

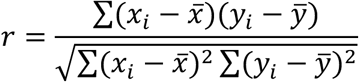

Where *x*_3_ and *y*_3_ are the values of observation *i* and ground truth *i*, respectively. *x^-^* and *y*z are the mean values of observation *i* and ground truth *i*.

### PSNR

Peak Signal-to-Noise Ratio (PSNR), which is the ratio of the maximum power of the signal to the noise power of the signal. It is a commonly used metric for evaluating image quality, which measures the quality of image recovery by comparing the distorted image with the original image. The higher the PSNR value, the smaller the error between the distorted image and the original image, and the better the image quality.

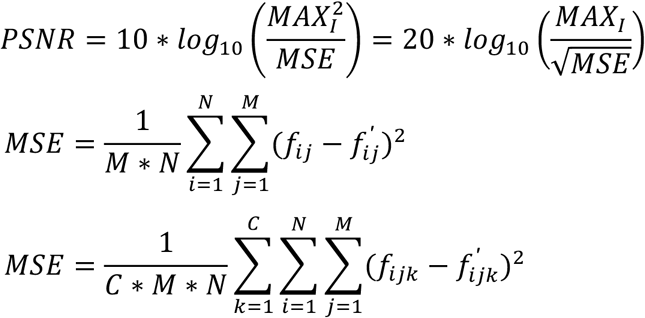

Where, *MAX*_I_ represents the maximum value of the pixel value in the image and *MSE* represents the mean value of the square of the difference between the corresponding pixels between two images.

### SSIM

Structural Similarity Index (SSIM) is a measure of the structural similarity of two images, which takes into account the brightness, contrast and structural information of the images. SSIM evaluates the image quality by comparing the local statistical properties of the images, which is closer to the human eye’s perception of the image quality. The higher the value of SSIM, the higher is the structural similarity of the two images. The higher the SSIM value, the higher the structural similarity between the two images and the better the image quality.

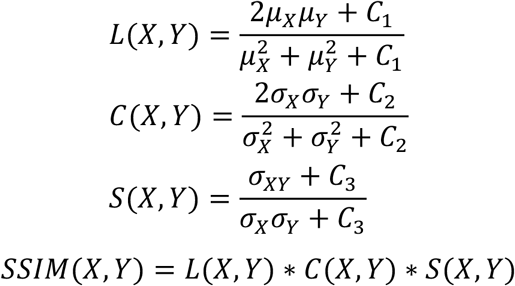

Where μ_x_ and μ_y_ are the mean values of the pixels of predicted *X* and ground truth *Y*, respectively. *x* and *y* are the standard values of the pixels of predicted *X* and ground truth *Y*, respectively. σ_x_ and σ_y_ represents the covariance of predicted *X* and ground truth *Y*, respectively. In addition, C_1_, C_2_, and C_3_ are constants.

### Nearest Neighbor Interpolation

Nearest Neighbor Interpolation **(NNI)**is the simplest method of up-sampling interpolation, i.e., making the value of the transformed pixel equal to the gray value of the input pixel nearest to it. The coordinates of the up-sampled high-resolution transcriptome matrix are calculated as

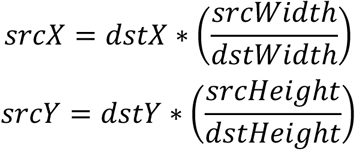

(*dstX*, *dstY*) is the horizontal and vertical coordinates of a pixel of the target image. *dstWidth* and *dstHeight* are the length and width of the target image.

*srcWidth* and *srcHeight* are the width and height of the original image. (*srcX*, *srcY*) is the coordinate of the original image corresponding to the target image at that point (*dstX*, *dstY*).

Each coordinate value *f*(*dstX*, *dstY*) of the high-resolution transcriptome matrix is calculated as:

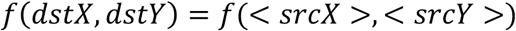

where<•>indicates rounding.

### Downstream missions

Normalization and principal components analysis (PCA) dimensionality reduction step was applied on each sample using *scanpy* Python package (version 1.10.2). Highly variable genes (HVGs) were identified using ‘highly_variable_genes’ function in *scanpy* Python package (version 1.10.2) and top 2048 HVGs were selected as input of STRESS. Spatial domain identification was achieved by default methods in scRNA-seq: Gaussian Mixture Modeling based clustering model using ‘GaussianMixture’ fuction in *sklearn* Python package (version 1.4.2), k-nearest neighbor graph based Leiden and Louvain clustering method in *scanpy* Python package (version 1.10.2).

Differentially expressed genes were identified using ‘rank_genes_groups’ in *scanpy* Python package (version 1.10.2). The gene set enrichment analysis (GSEA) was performed using ‘enrichr’ function in gseapy Python package (version 1.1.6) and reference gene sets KEGG and GO were load through ‘MyGeneInfo’ fuction in mygene Python package (version 3.2.2).

### Clustering evaluation metrics

Clustering performance was evaluated using 4 metrices: Normalized Mutual Information (NMI), Adjusted Rand Index (ARI), Silhouette Coefficient (SC) and Davies-Bouldin Index (DBI). All metrics were calculated using functions ‘normalized_mutual_info_score’, ‘adjusted_rand_score’, ‘silhouette_score’, and ‘davies_bouldin_score’ from sklearn Python package (version 1.4.2).

## Data availability

**Human dorsolateral prefrontal cortex dataset** (DLPFC) generated from 10X Visium platform. Raw data including 12 different tissue slices was downloaded from http://spatial.libd.org/spatialLIBD/. **Human HER2-positive breast cancer dataset** (HER2ST) generated from Spatial Transcriptomics (ST) platform. Raw data including 36 tissue slices from 8 samples was downloaded from https://github.com/almaan/her2st. **Mouse Organogenesis Spatiotemporal Transcriptomic Atlas** (MOSTA) generated from Stereo-seq platform. Raw data including 8 tissue slices was downloaded from https://db.cngb.org/search/project/CNP0001543. **Mouse hippocampus dataset** generated from Slide-seqV2 platform was used to simulate 10X Visium data. Raw data including sample ‘Puck_200115_08’ was downloaded from https://singlecell.broadinstitute.org/single_cell/study/SCP815/sensitive-spatial-genome-wideexpression-profiling-at-cellular-resolution#study-summary. **Mouse brain serial section dataset** generated from 10X Visium platform. Raw data including 4 serial slices from 2 sections was downloaded from https://www.10xgenomics.com/datasets/mouse-brain-serial-section-1-sagittal-anterior-1-standard-1-0-0, https://www.10xgenomics.com/datasets/mouse-brain-serial-section-1-sagittal-posterior-1-standard-1-0-0, https://www.10xgenomics.com/datasets/mouse-brain-serial-section-2-sagittal-anterior-1-standard-1-0-0 and https://www.10xgenomics.com/datasets/mouse-brain-serial-section-2-sagittal-posterior-1-standard-1-0-0. **Human invasive ductal carcinoma dataset** generated from 10X Visium platform. Raw data including 1 slice collected from IDC patient was downloaded from 10X dataset https://www.10xgenomics.com/datasets/invasive-ductal-carcinoma-stained-with-fluorescent-cd-3-antibody-1-standard-1-2-0. **Visium HD dataset** generated from 10X Visium HD platform. Raw data including 6 different samples was downloaded from 10X dataset website: Human Colorectal Cancer (FFPE) sample from https://www.10xgenomics.com/datasets/visium-hd-cytassist-gene-expression-libraries-of-human-crc, Human Lung Cancer (FFPE) sample from https://www.10xgenomics.com/datasets/visium-hd-cytassist-gene-expression-human-lung-cancer-post-xenium-expt, Human Breast Cancer (FFPE) sample from https://www.10xgenomics.com/datasets/visium-hd-cytassist-gene-expression-libraries-human-breast-cancer-ffpe-if, Human Breast Cancer (Fresh Frozen) sample from https://www.10xgenomics.com/datasets/visium-hd-cytassist-gene-expression-libraries-human-breast-cancer-ff-ultima, Human Kidney (FFPE) sample from https://www.10xgenomics.com/datasets/visium-hd-cytassist-gene-expression-libraries-human-kidney-ffpe, Human Tonsil (Fresh Frozen) sample from https://www.10xgenomics.com/datasets/visium-hd-cytassist-gene-expression-libraries-human-tonsil-ff-ultima.

## Code availability

STRESS is implemented based on python 3.8.0. Other tools and packages used in the data analysis includes: numpy 1.24.1, einops 0.7.0, jupyter 1.1.1, leidenalg 0.10.2, Louvain 0.8.2, mamba-ssm 1.2.0.post1, pandas 2.0.3, scanpy 1.9.6, scikit-learn 1.3.2, scipy 1.10.1, seaborn 0.13.2, torch 2.1.1+cu118, torchaudio 2.1.1+cu118, torchmetrics 1.4.0.post0, torchvision 0.16.1+cu118, transformers 4.41.2, umap-learn 0.5.5. Source codes in this study will be maintained and updated at https://github.com/YYYYYeFei/STRESS.git.

## Acknowledgements

This work is supported by Scientific Research Start-up Funds (QD2021005N), Shenzhen Science and Technology Program (WDZC20220819134430002) and Shenzhen Universities Stable Funding Key Projects (WDZC20200821104802001).

