## Supplementary Figure 1-14 for "STRESS: Spatial Transcriptome Resolution Enhancing Method based on the State Space Model"

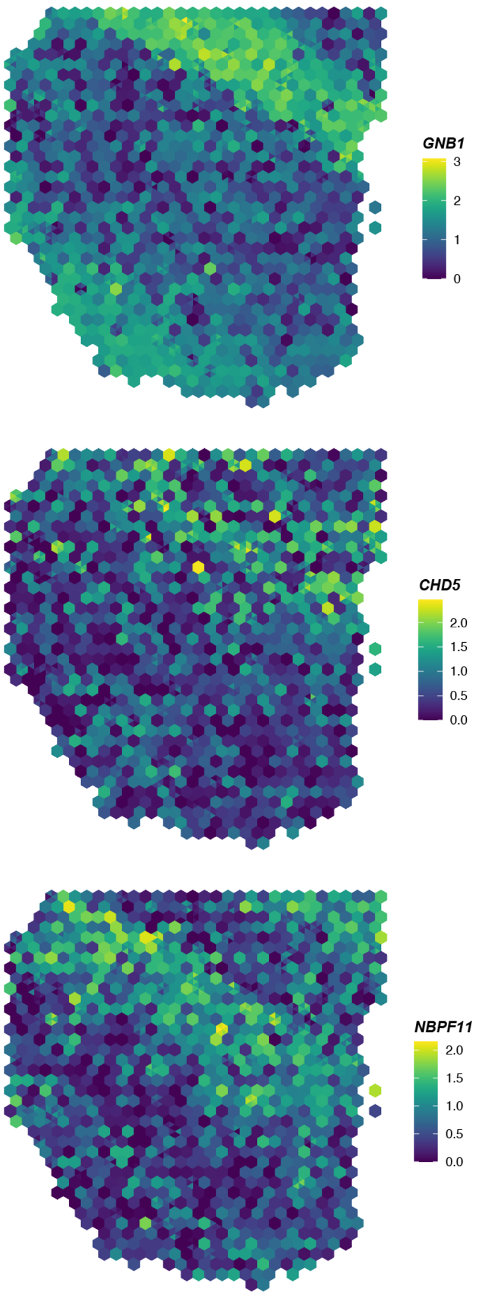


**Supplementary Fig.1: Genes expression distributions in DLPFC predicted by BayesSpace**

Spatial distributions of gene GNB1, CHD5 and NBPF11 in sample 151507 in DLPFC dataset predicted by BayesSpace using low resolution data as input.

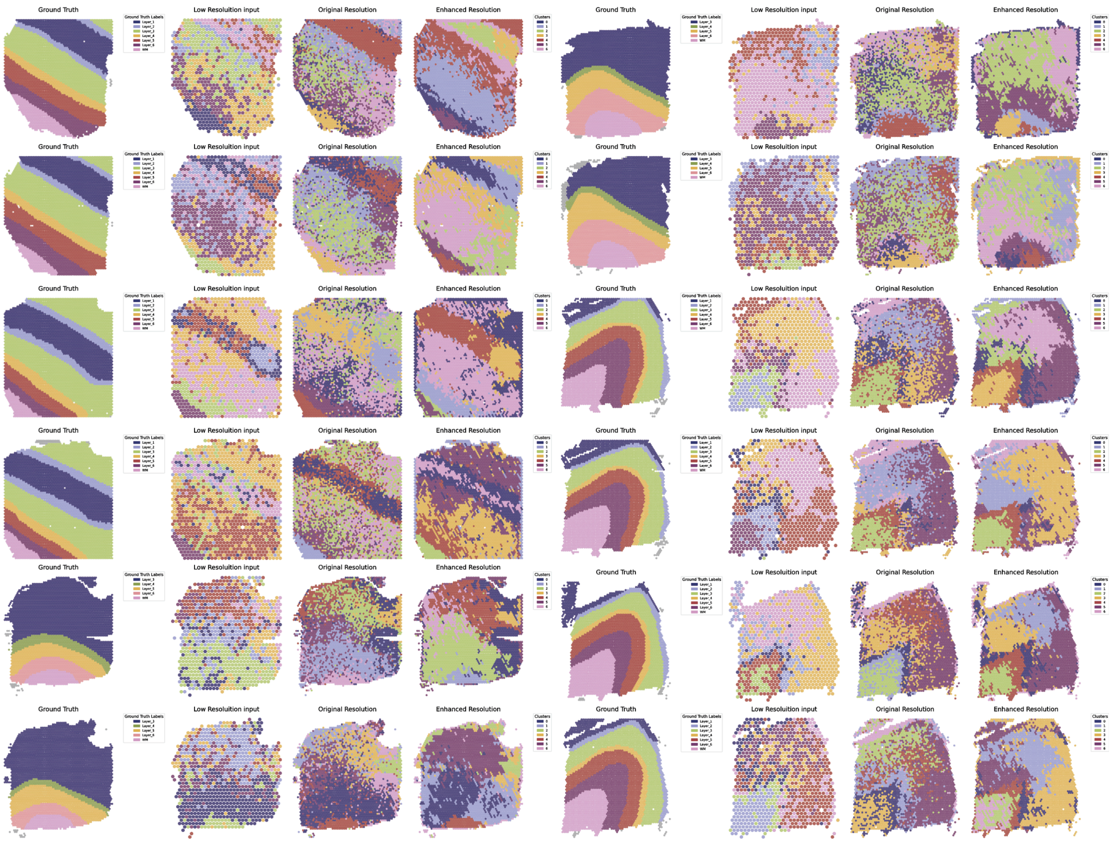


**Supplementary Fig.2: Clustering results of all samples in DLPFC dataset**

Manual annotations of all samples in DLPFC dataset, spatial domain identification results based on low resolution input, original resolution Ground Truth and STRESS-enhanced resolution prediction using same clustering method and same parameters

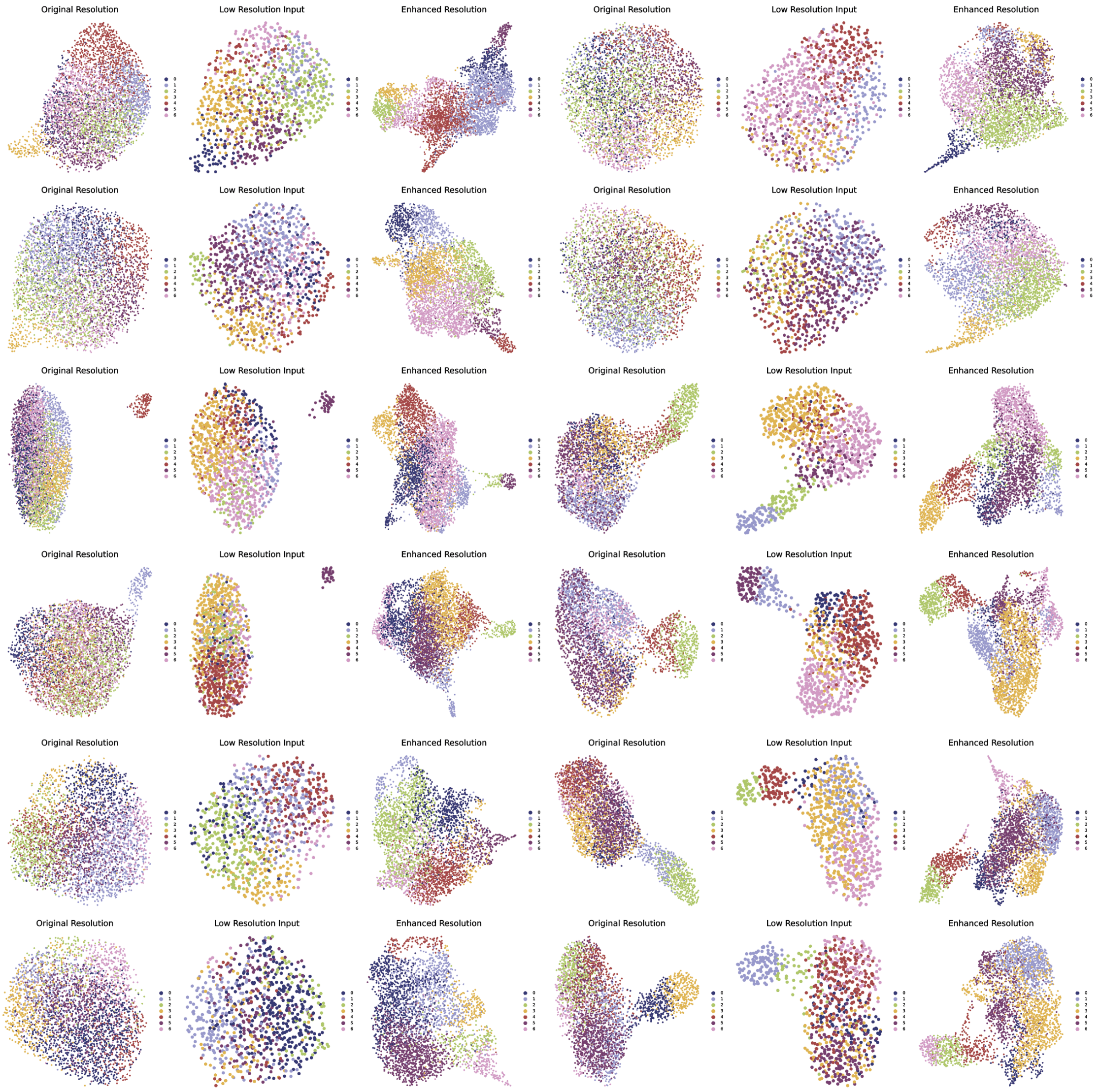


**Supplementary Fig.3: Latent representations of all samples in DLPFC dataset**

Latent representations of all samples in DLPFC dataset obtained by performing UMAP dimensionality reduction on gene expression matrices of low-resolution input, original resolution Ground Truth, STRESS-Visium-enhanced resolution prediction, with different colors representing previously identified domain labels

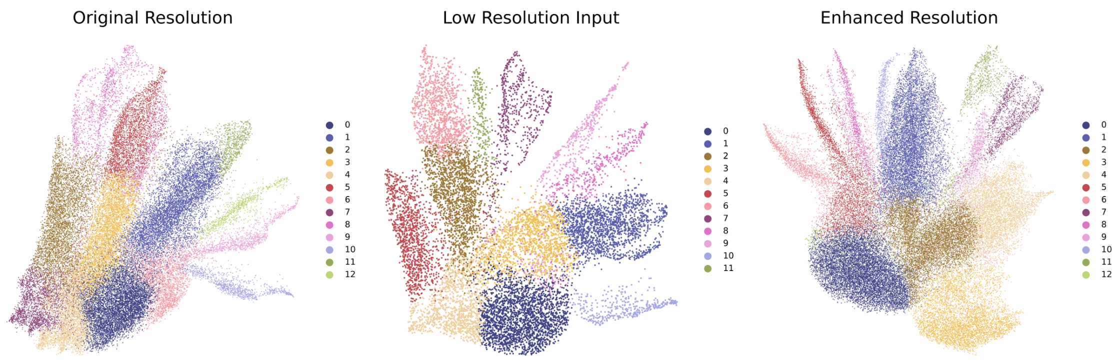


**Supplementary Fig.4: Latent representations of Slide-seq V2 dataset**

Latent representations of the mouse sample in Slide-seq V2 dataset obtained by performing UMAP dimensionality reduction on gene expression matrices of low-resolution input, original resolution Ground Truth, STRESS-Visium-enhanced resolution prediction, with different colors representing previously identified domain labels

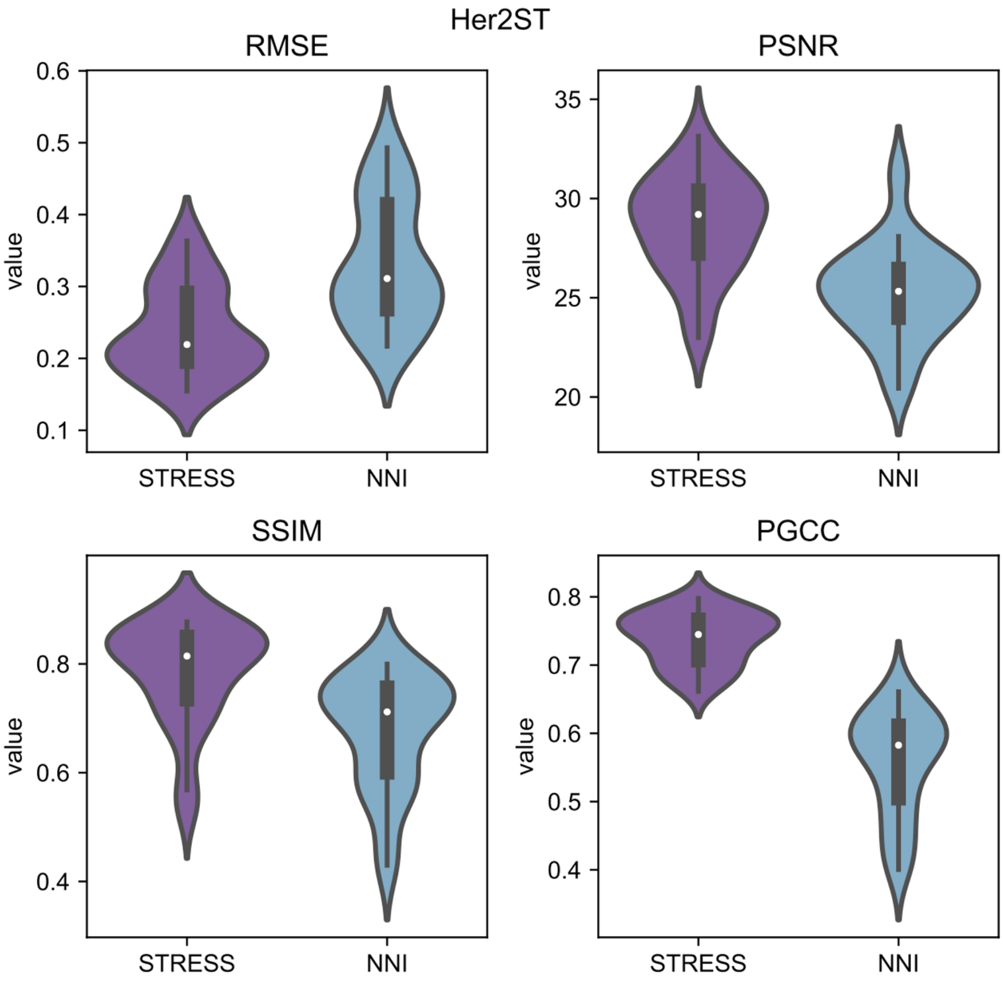


**Supplementary Fig.5: STRESS-ST performance compared to NNI method.** RMSE, Root Mean Square Error; PSNR, Peak Signal-to-Noise Ratio; SSIM, Structural Similarity Index; PGCC, per-gene expression correlation coefficient; NNI, Nearest Neighbor Interpolation.

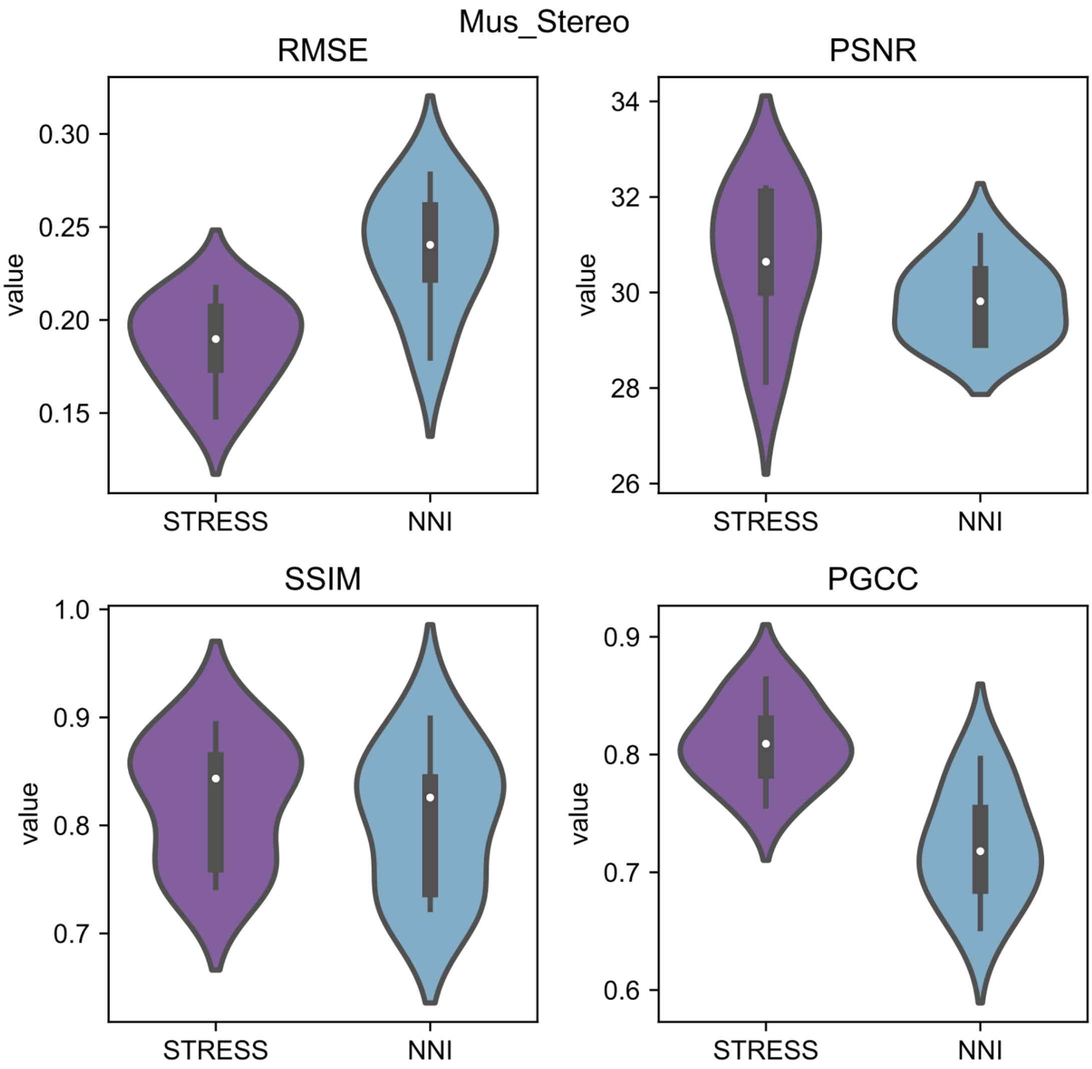


**Supplementary Fig.6: STRESS-Stereo performance compared to NNI method.** RMSE, Root Mean Square Error; PSNR, Peak Signal-to-Noise Ratio; SSIM, Structural Similarity Index; PGCC, per-gene expression correlation coefficient; NNI, Nearest Neighbor Interpolation.

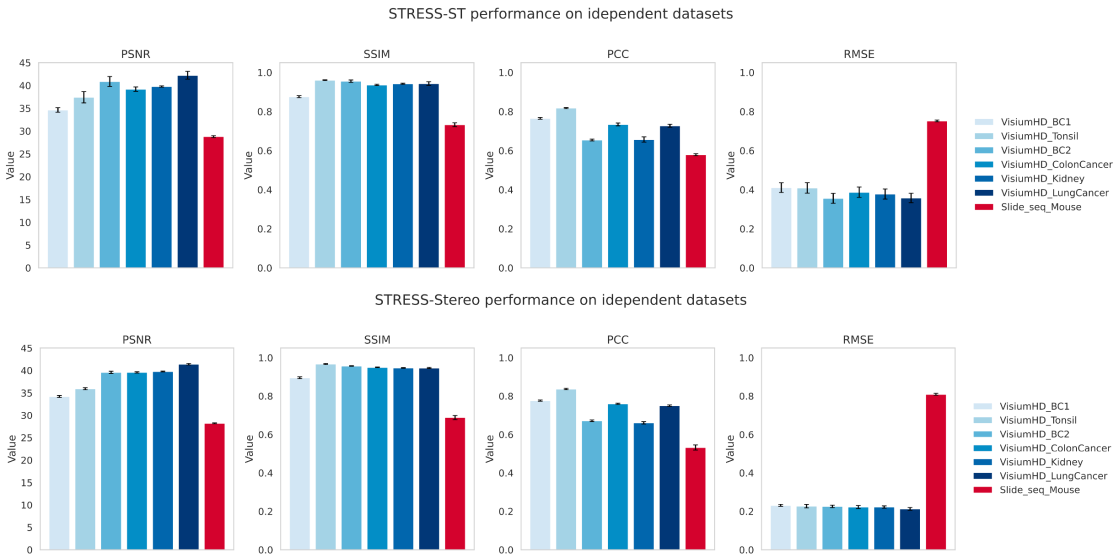


**Supplementary Fig.7: STRESS-ST and STRESS-Stereo performance on different samples of independent datasets (Visium HD and Slide-seq V2).** RMSE, Root Mean Square Error; PSNR, Peak Signal-to-Noise Ratio; SSIM, Structural Similarity Index; PCC, per-gene expression correlation coefficient; NNI, Nearest Neighbor Interpolation.

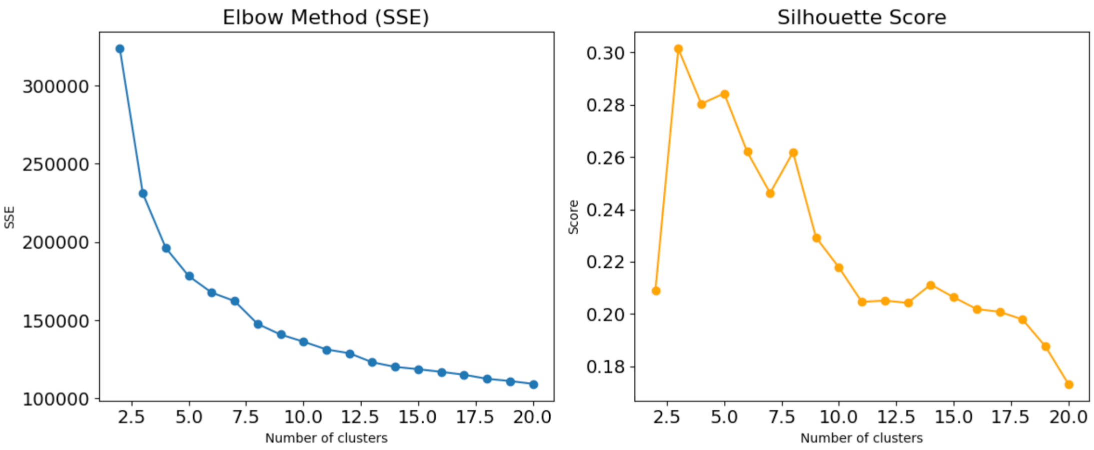


**Supplementary Fig.8: Optimal cluster number estimation.**

The elbow plot (left) shows the decrease in SSE (Sum of Squared Errors) with increasing the cluster number, while the silhouette score plot (right) evaluates clustering quality. Both metrics guide the selection of the optimal number of clusters.

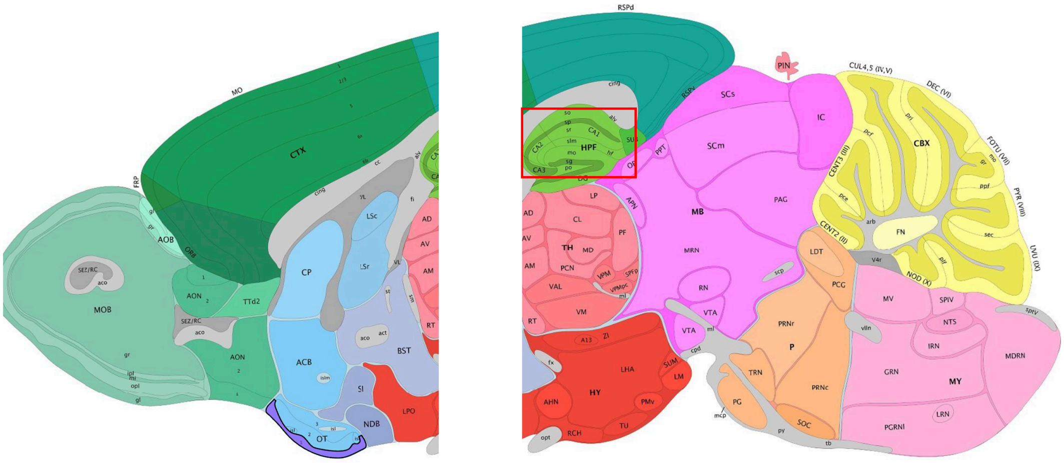


**Supplementary Fig.9: The Allen Brain Atlas[23,36] annotated layer structure of mouse brain posterior and anterior.**

Red Box, HPF region, Hippocampal formation; CBX, Cerebellum cortex; gr, granular layer; MY, Medulla; P, Pons; MB, Midbrain; TH, Thalamus; HY, Hypothalamus; BST, Bed nuclei of the stria terminalis; SI, Substantia innominate; NDB, Diagonal band nucleus; CP, Caudoputamen; ACB, Nucleus accumbens; CTX, Cerebral cortex; OT, Olfactory tubercle; CLA, Claustrum; AON, Anterior olfactory nucleus; PIR, Piriform area; AOB, Accessory olfactory bulb; MOB, Main olfactory bulb.

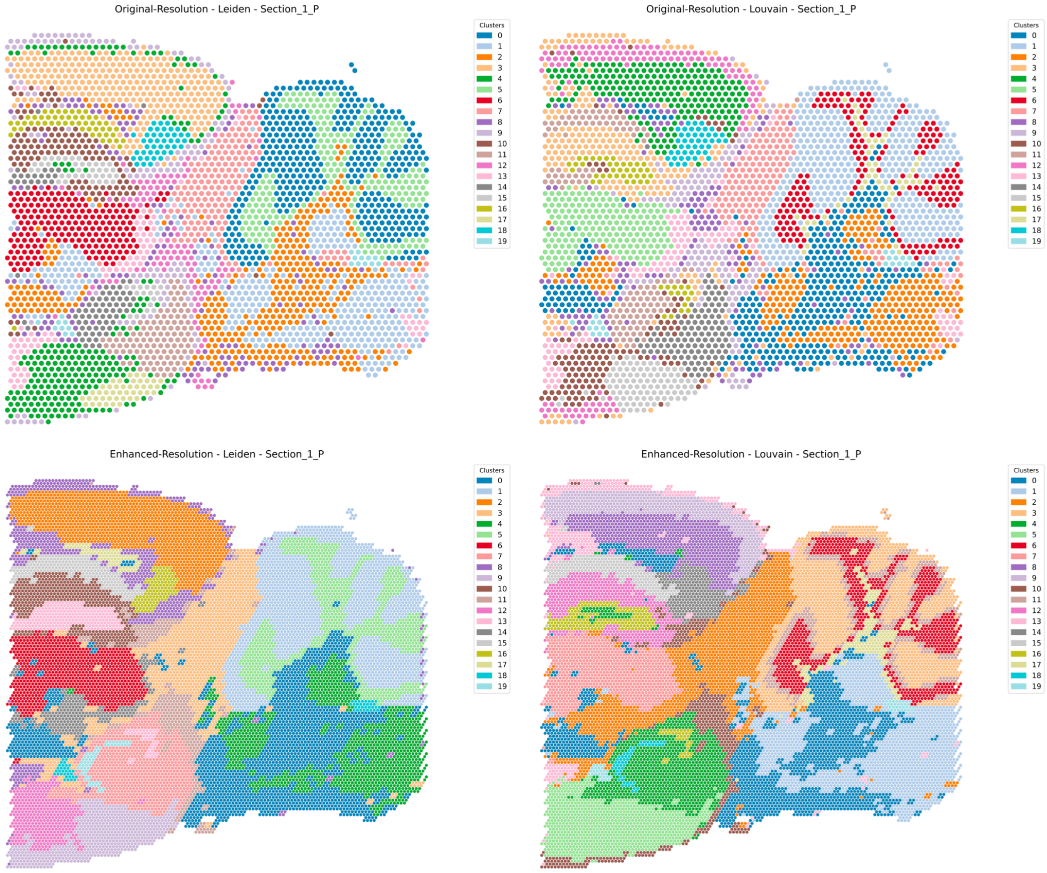


**Supplementary Fig.10: Spatial domain identification results with increased cluster numbers.**

Spatial domain identification results of mouse brain posterior section 1 using two different clustering methods (Leiden and Louvain) at original and STRESS-enhanced resolutions.

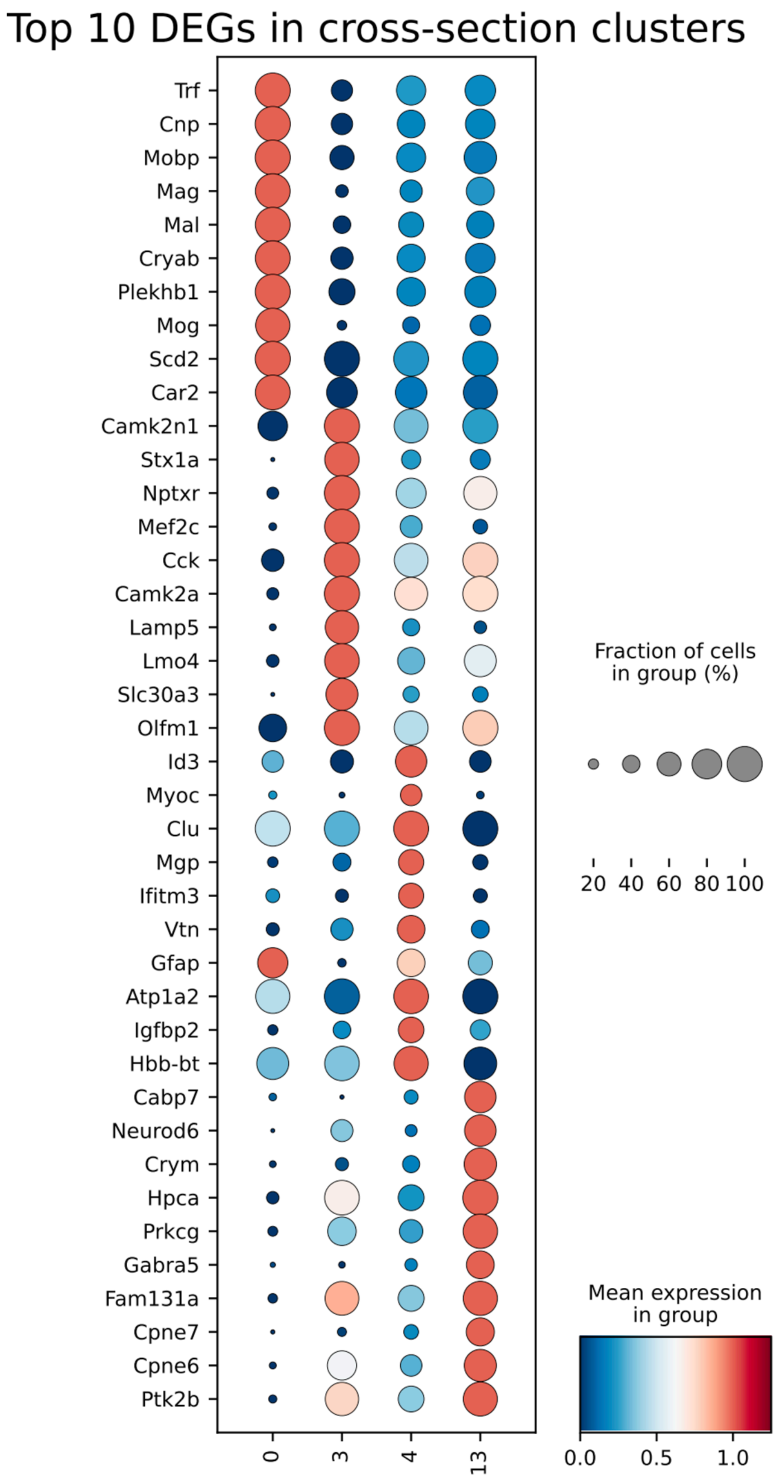


**Supplementary Fig.11: DEGs in cross-section clusters in Mouse Brain dataset**

Top 10 differential expression genes among cluster 0, 3, 4, 13 in integrated mouse brain sample at STRESS-enhanced resolution.

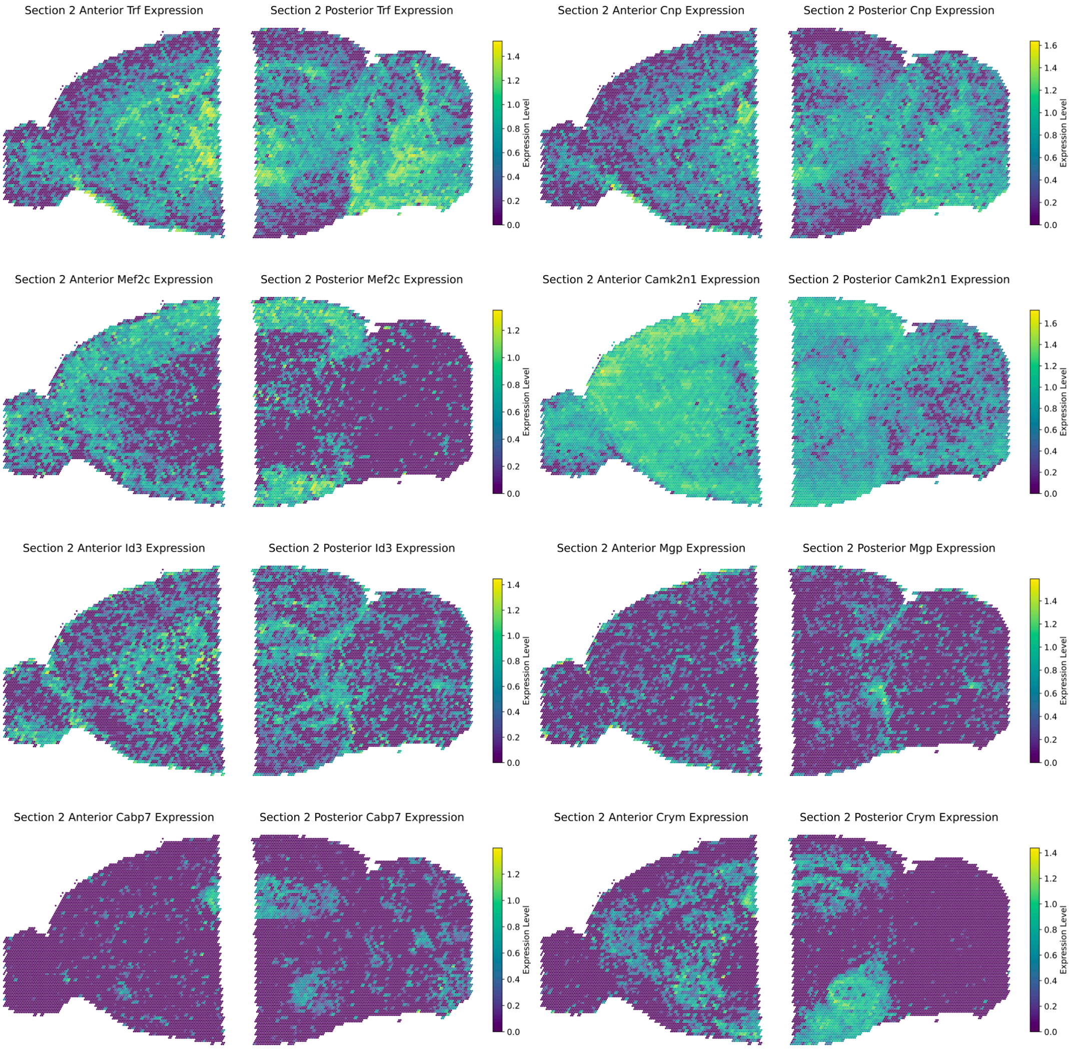


**Supplementary Fig.12: Genes expression distributions of two adjacent samples in mouse brain section 2.**

Distributions of genes enriched in different clusters. *Trf* and *Cnp* in cluster 0, *Mef2c* and *Camk2n1* in cluster 3, *Id3* and *Mgp* in cluster 4, *Cabp7* and *Crym* in cluster 13.

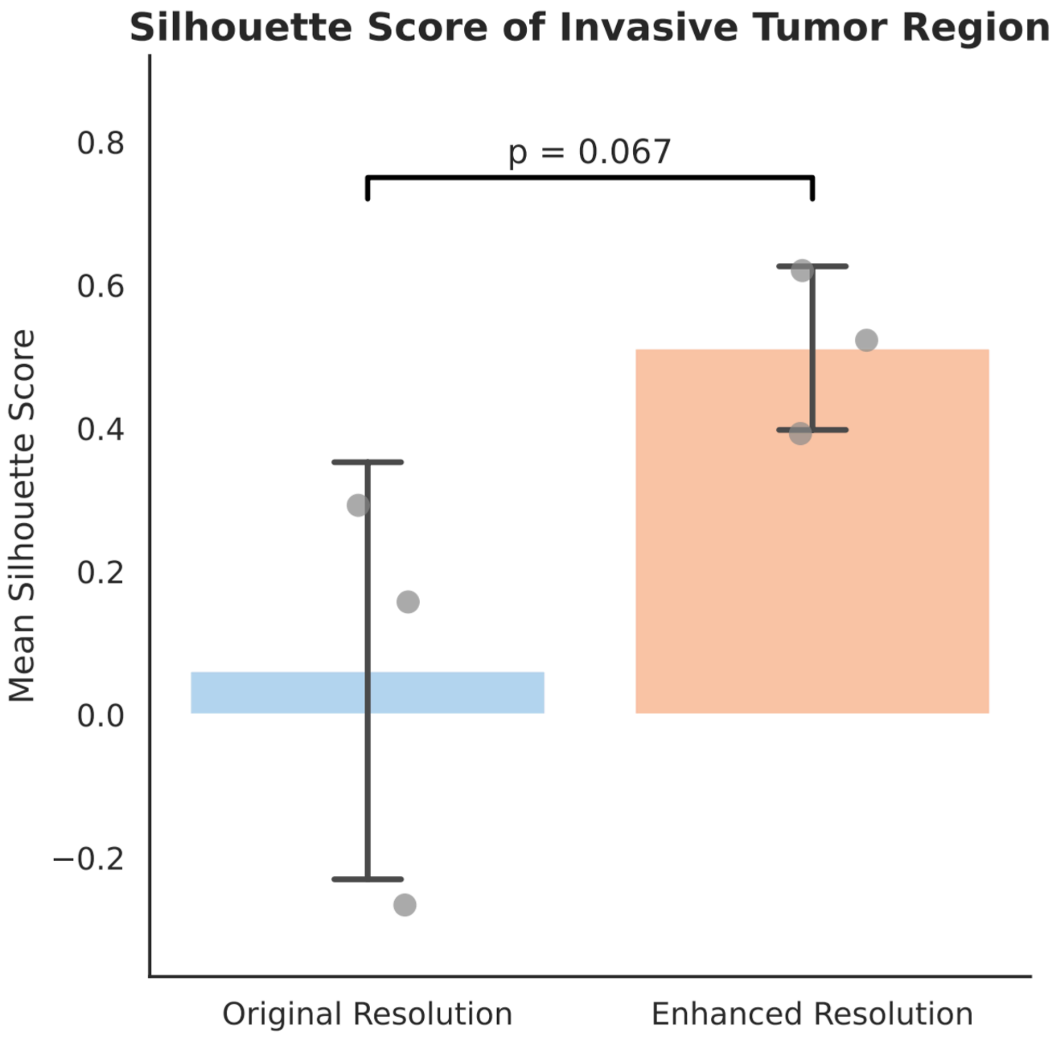


**Supplementary Fig.13: Comparison of silhouette scores between original and enhanced resolution clustering.**

Bar plot showing the mean silhouette scores of selected clusters (three per condition) in the invasive tumor region under original and enhanced resolution. Each point represents the average silhouette score of a single cluster. Enhanced resolution significantly improved clustering quality ($p$ = 0.067, two-sided $t$-test).

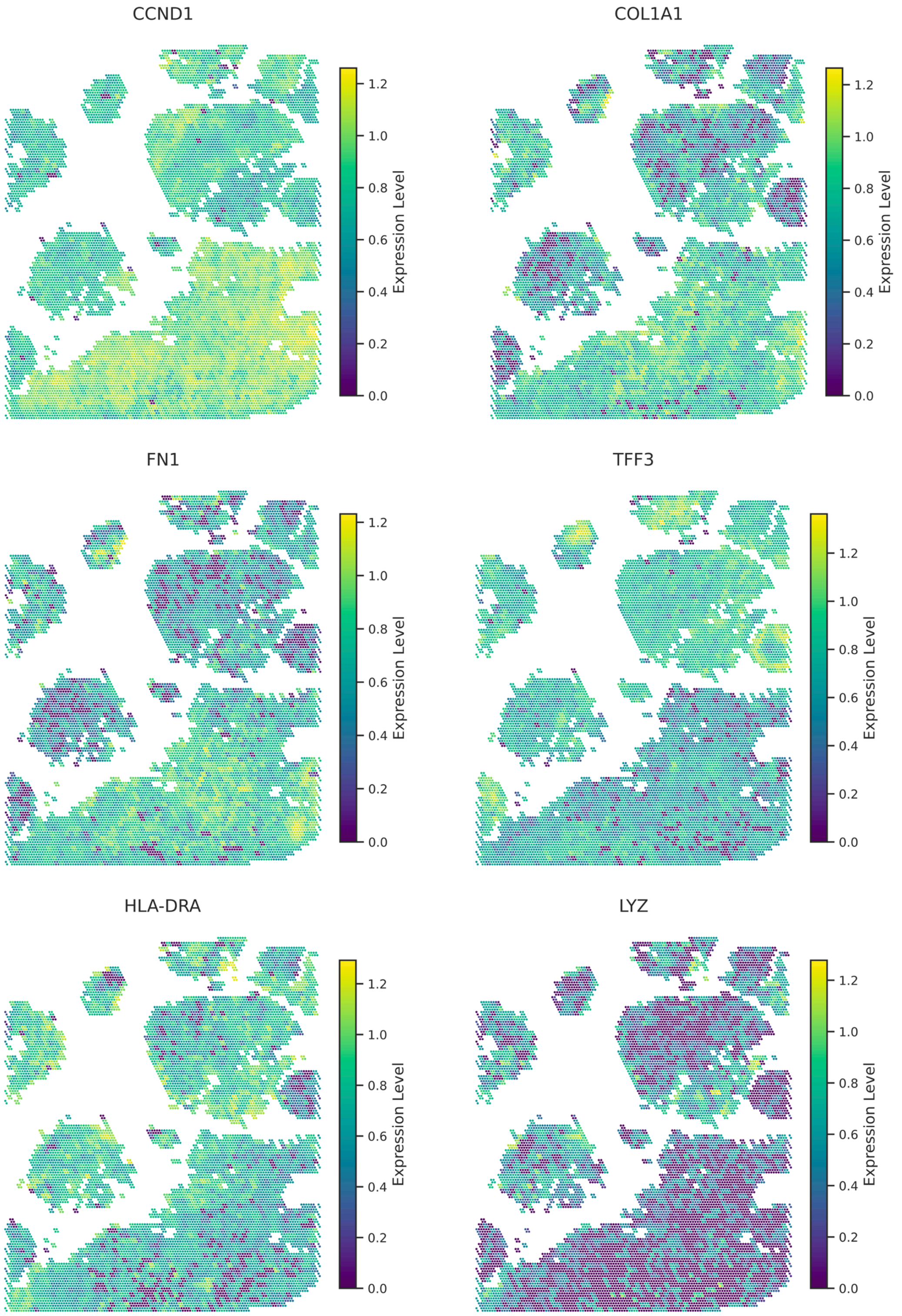


**Supplementary Fig.14: Genes expression distributions of tumor tissue regions in the IDC patient.**

Distributions of genes enriched in different spatial regions. *CCND1*, *COL1A1* and *FN1* in invasive carcinoma region, *TFF3* in carcinoma in situ region, *HLA-DRA* and *LYZ* in benign hyperplasia region.
